# Serum-dependent and independent regulation of PARP2

**DOI:** 10.1101/483289

**Authors:** Qizhi Sun, Mohamed I. Gatie, Gregory M. Kelly

**Author notes:** Address correspondence to: G.M. Kelly, 1151 Richmond St. London, ON Canada; To whom correspondence should be addressed.

## Abstract

PARP2 belongs to a family of proteins involved in cell differentiation, DNA damage repair, cellular energy expenditure, chromatin modeling and cell differentiation. In addition to these overlapping functions with PARP1, PARP2 participates in spermatogenesis, T-cell maturation, extraembryonic endoderm formation and adipogenesis. The function(s) of PARP2 is far from complete, and the mechanism(s) by which the gene and protein are regulated are unknown. In this study, we found that two different mechanisms are used *in vitro* to regulate PARP2 levels. In the presence of serum, PARP2 is degraded through the ubiquitin-proteasome pathway, however, when serum is removed, PARP2 is rapidly sequestered into an SDS- and urea-insoluble fraction. This sequestration is relieved by serum in a dose-dependent manner, and again PARP2 is detected by immunoblotting. Furthermore, and despite the presence of a putative serum response element in the *PARP2* gene, transcription is not affected by serum deprivation. These observations that PARP2 is tightly regulated by distinct pathways highlights the critical roles PARP2 plays under different physiological conditions.

## Introduction

The poly-ADP-ribose polymerase (PARP) enzymes belong to a large family of proteins, including 17 in humans (Leung 2014), that have integral roles in DNA repair, and thus are studied extensively as targets to inhibit a variety of cancers (Ali et al. 2016). PARPs are also involved in the inflammatory response, transcription, mitochondrial function, oxidative metabolism and heterochromatin function (Ali et al. 2016; Bai and Canto 2012; Chen et al. 2018; Dantzer and Santoro 2013; Gupte et al. 2017; Hottiger 2015; Jeggo 1998; Krishnakumar and Kraus 2010). PARPs catalyze the formation of free poly-ADP-ribose polymers as well as poly-ADP-ribosylated (pARylated) proteins using NAD^+^ as a substrate (Kraus 2015; Nicolas et al. 2010). PARP1 and PARP2 share similar crystal structure and function (Hanzlikova et al. 2017; Oliver et al. 2004), and both proteins homo-and heterodimerize and poly ADP-ribosylate each other (Schreiber et al. 2002). Despite these structural and functional similarities, PARP2 can function in a manner distinct from PARP1 (Bai et al. 2011; Celik-Ozenci and Tasatargil 2013; Dantzer et al. 2006b; Farres et al. 2015; Jha et al. 2009; Kamboj et al. 2013; Nicolas et al. 2010; Tramontano et al. 2007; Yelamos et al. 2006), but both proteins are required for mouse early development (Menissier de Murcia et al. 2003; Nicolas et al. 2010). This early requirement is evident from studies with mouse F9 teratocarcinoma cells, where the loss of PARP1 or PARP2 blocks the expression of markers (Quenet et al. 2008) required in the Wnt- and Hedgehog-dependent differentiation of extraembryonic endoderm (Deol et al. 2017; Golenia et al. 2017; Hwang and Kelly 2012). Another important role assigned to PARP proteins is their maintenance of telomere integrity (Dantzer et al. 2004), and genome stability through recruiting DNA repair factors to DNA-strand breaks and base-excision lesions resulting from DNA damage (Ame et al. 1999; Riccio et al. 2016; Schreiber et al. 2002). These activities are suspended during apoptosis by caspase-8, which serves to inactivate PARP2 (Benchoua et al. 2002). In addition, PARP2 contributes to the translocation of apoptosis-inducing factor (AIF) from the mitochondria to the nucleus during oxidative damage to DNA (Li et al. 2010), and it can control cell death and autophagy linked to oxidative stress (Wyrsch et al. 2012). PARP2, together with the PPARγ/RXR transcription machinery, is also important in adipocyte differentiation (Bai et al. 2007), in the regulation of surfactant protein B expression in pulmonary cells (Maeda et al. 2006), and for hematopoietic stem cell survival under steady-state conditions and in response to radiation stress (Farres et al. 2013). However, the loss of PARP2 can increase apoptosis, contradicting a focal cerebral ischemia report that showed a suppression of AIF in PARP2^-/-^ (Li et al. 2010). Despite these discrepancies, owing to cell specificity and exemplified by the PARP2 depletion results in several cell lines (Boudra et al. 2015), these studies confirm that PARP2 is not entirely functionally redundant with PARP1. Furthermore, they underscore the importance of PARP2 in maintaining a number of key cellular physiological processes. For these reasons, we sought to investigate the mechanism by which PARP2 is regulated under normal and stress-induced conditions.

To investigate how PARP2 is regulated, a survey of several cell lines showed robust PARP2 levels that were stable over several hours of cycloheximide treatment. Interestingly, following serum removal, PARP2 signals were absent, but when the cells were returned to complete medium, the protein was detected on immunoblots. Since *PARP2* gene expression was not affected by serum removal, we postulated the loss of PARP2 was the result of post-translational modifications. Analysis using proteolytic inhibitors failed to identify the protease(s) responsible for the loss of PARP2 under serum-free conditions. Finally, our focus turned to the proteasome and the post-translational modification of PARP2 by ubiquitination. *In vitro* assays showed that PARP2 was ubiquitinated and when cells were cultured in complete medium, ubiquitination led to PARP2 degradation in the proteasome. Unexpectedly, inhibiting proteasome activity under serum-free conditions did not prevent PARP2 signals from disappearing, which together with the protease inhibition experiments, suggested the protein was being sequestered to a denaturing-insoluble compartment rather than being degraded by a protease or the proteasome.

Together, our findings strongly support the notion that when serum is present, and cells are stimulated to grow, PARP2 is detected under standard SDS-denaturing conditions and thus is available fulfill its roles in maintaining normal cellular physiology. However, in the absence of serum, PARP2 is sequestered to an SDS-and urea-resistant compartment. Regardless of the mechanism, this outcome would serve to reduce global ADP-ribosylation enzyme activity to thereby minimize energy expenditure under adverse conditions.

## Materials and methods

### Antibodies, plasmids and reagents

PARP1/2 (H250), β-actin and ERK antibodies were purchased from Santa Cruz Biotechnology, rabbit anti-mouse PARP2 (Yucatan) from Enzo Life Sciences, Inc., rabbit anti-human PARP2 antibody from Axxora, and GST and HA antibodies from GenScript. HRP-conjugated secondary antibodies were purchased from Pierce. *pBC-GST-PARP2*, *pBC-GST-NLS* and *pEGFP-PARP2* plasmids were gifts of Dr. V. Schreiber (École Supérieure de Biotechnologie Strasbourg, France). The *mRFP-ub* plasmid was a gift of Dr. N. Dantuma (Karolinska Institutet, Sweden; Addgene #11935), and the *pMT123-HA-ubiquitin* plasmid was kindly provided by Dr. D. Bohmann (University of Rochester, USA). MG-132 and cycloheximide were from Sigma, and the caspase inhibitor III (BD-FMK), calpeptin and pepstatin A methyl ester (PME) were from Calbiochem (EMD Millipore). Caspase-8 inhibitor (Z-IETD-FMK) was from BD-Biosciences. Alpha-2-macroglobulin was purchased from Enzo Life Sciences and leupeptin from Bio Basic Inc. The HALT protease inhibitor cocktail was from Pierce and GST-PARP2 human recombinant protein from BPS Bioscience. Protein fraction II, HA-ubiquitin, ubiquitin aldehyde and ubiquitin conjugation reaction buffer kits were purchased from Boston Biochem, Inc. The transfection reagent XtremeGene 9 was from Roche Applied Sciences, and Glutathione Sepharose 4B beads were purchased from GE Healthcare Life Sciences. Power SYBR Green PCR master mix was purchased from Invitrogen Thermo Fisher Scientific.

### Cell culture, treatment and transfection

COS-7, MCF-7, HeLa, NIH3T3, MEF F20 and IMCD cells were maintained in Dulbecco’s modified Eagle’s medium (DMEM)/F-12 or DMEM supplemented with 10% FBS, 100 units/ml penicillin and 100 mg/ml streptomycin in 5% CO2 at 37°C. Cells were treated with different protease inhibitors at the concentration and duration as indicated in the figures. Cells were subject to serum starvation or maintained in medium containing different concentrations of sera where stated. Transfections were carried out using X-tremeGENE 9 transfection reagent as per manufacturer’s recommendation. To test if the loss of the PARP2 signal on blots under serum starvation conditions was due to sequestration rather than degradation, MCF-7 cells were cultured to approximately 90% confluence in complete medium (CM) and then this was removed and replaced with serum-free (SF) medium containing 50µgml^-1^ cycloheximide (CHX) to inhibit protein synthesis. The cells were incubated for 7hr, and then the medium replaced with CM containing 50µgml^-1^ CHX. The cells were cultured for 1hr and then lysed in Laemmli sample buffer. Cells serum-starved for 7hr with or without CHX treatment or remaining in CM with or without CHX.

### End point and quantitative reverse transcription-PCR

MCF-7 cells were cultured in CM until 90% confluence. At this point the medium was removed for one plate and replaced with SF medium. Cells on a second plate were maintained in CM. Following 30 minutes, cells on both plates were lysed in TRIzol (Invitrogen Thermo Fisher Scientific) and total RNA extracted following the manufacturer’s instructions. Following preparation of first strand cDNA by reverse transcription with (+) or without reverse transcriptase (-) (control), PCR was performed using primers specific to human *PARP2* (forward 5′- GAATCTGTGAAGGCCTTGCTG- 3′ and reverse 5′-TTCCCACCCAGTTACTCATCC-3′). PCR products were resolved on 1% agarose gels and images captured with a FluorChem IS-8900 Imager (Alpha Innotech Corp.). For q-RT-PCR, MCF-7 cells were cultured in CM, or serum-starved for 15, 30 and 60 minutes. F9 cells were also serum-starved for 60 minutes and then treated for additional 12 hours with DMSO, or with retinoic acid (RA) 10^−7^ M. For controls, cells were treated for 12 hours in CM containing DMSO or RA 10^−7^ M. In all experiments, total RNA was extracted with the RNeasy mini kit (QIAGEN), and first strand cDNA prepared using qScript cDNA SuperMix (Quanta Bioscience) following the manufacturer’s instructions. For MCF-7 cells q-RT-PCR was performed using the abovementioned *PARP2* primers, while mouse *Gata6* primers (forward 5′-ATGGCG TAGAAATGCTGAGG-3′ and reverse 5′-TGAGGTGGTCGCTTGTGTAG-3′) and *Hoxb1* primers (forward 5′- GGGGTCGGAATCTAGTCTCCC-3′ and reverse 5′-CCTCCAAAGTAGCCATAAGGCA) were used with the F9 cells. *GAPDH* primers (forward 5′-GTGTTCCTACCCCCAATGTGT-3′ and reverse 5′- ATTGTCATACCAGGAAATGAGCTT-3′) were used as the reference primers for MCF-7 cells and mouse *L14* (forward 5′-GGGTGGCCTACATTTCCTTCG-3′ and reverse 5′- GAGTACAGGGTCCATCCACTAAA-3′) for F9 cells. Reactions were performed using a CFX Connect Real-Time PCR Detection System (Bio-Rad), and results were analyzed using the comparative cycle threshold (2^-ΔΔCt^) method with the internal controls.

### Cell free ubiquitination and GST pull-down assays

COS-7 cells were transfected with *pBC-GST-PARP2* or *pBC-GST-NLS* (control) alone or with *pMT123*-*HA-ubiquitin* plasmids using the X-tremeGENE 9 transfection reagent (Sigma-Aldrich). Twenty-four hours post-transfection, one plate of COS-7 cells transfected with both *pBC-GST-PARP2* and *pMT123-HA-ubiquitin* was treated with 40μM MG-132, and the second with DMSO as a vehicle control. After 20hr, cells were washed in PBS (phosphate buffered saline) and proteins harvested by lysing in 1x RIPA buffer (50mM Tris-HCl pH 7.5, 150mM NaCl, 1% NP-40, 0.5% Na-deoxycholate and 0.1% SDS) supplemented with protease inhibitor cocktail on ice. Cell lysates were stored at -80°C for GST-pull down assays. Protein concentrations were determined using a Bradford assay and equal amounts of total protein from each sample were incubated with Glutathione Sepharose 4B beads overnight at 4°C. Beads were washed 4 × 5 minutes with RIPA buffer and then resuspended in 2x Laemmli sample buffer. Proteins pull-downed in these assays were resolved by 8% SDS-PAGE, and then subjected to immunoblot analysis.

### *In vitro* ubiquitination assay

GST-PARP2 human recombinant protein (0.4μg) was added to a 20μl final volume reaction mix containing 1x ubiquitin conjugation reaction buffer, 0.5mM MG-132, 1x ubiquitin aldehyde, 2mM HA-ubiquitin, 1x Mg-ATP and 1μl HeLa protein fraction II. For controls, 1μl of water was substituted for 1μl HeLa protein fraction II. Reactions were carried out at 37°C for 2hr and then inhibited by adding 2μl 10x E1 stopping buffer, 4μl 5x Laemmli sample buffer and 1.5μl of β-mercaptoethanol. Samples were boiled for 5 minutes, and the proteins resolved by 8% SDS-PAGE before immunoblot analysis.

### Immunoblotting

After a PBS wash, cells were either lysed on ice in 1x Laemmli sample buffer supplemented, or urea lysis buffer (8M Urea, 2M Thiourea, 4% w/v CHAPS). All buffers were supplemented with protease inhibitor cocktail, but without bromophenol blue. Lysates were sonicated for 10sec and boiled for 5 minutes, then centrifuged for 10 minutes at 13,200 rpm at 4°C. To determine protein concentrations using a Bradford assay, aliquots were diluted 200 fold to minimize SDS interference. Equal amount of proteins was resolved by 8% or 10% SDS-PAGE and then transferred to nitrocellulose membranes, which were then blocked in 5% skim milk/Tris-buffered saline/Tween 20 (TBS/T) buffer for 2hr at room temperature. Following an overnight incubation in primary antibody (1:2500 diluted in blocking buffer) at 4°C, membranes were washed in TBS/T buffer and then incubated 1hr with a HRP-conjugated secondary antibody (1:4000 diluted in blocking buffer) at room temperature. Membranes were washed extensively with TBS/T buffer and signals were detected by enhanced chemiluminescence (Pierce Thermo Fisher Scientific). Densitometric analyses were performed using ImageJ software (NIH).

### Confocal microscopy

HeLa cells cultured on glass cover slips were transfected with *pEGFP-PARP2* and m*RFP-ub* plasmids. At 24hr post-transfection, cells were fixed with 4% paraformaldehyde in PBS for 30 minutes at room temperature. After 3 × 10 minute washes with PBS, cells were mounted in ProLong Gold anti-fade mounting medium (Invitrogen Thermo Fisher Scientific) and viewed with a Zeiss LSM 510 Duo Vario confocal microscope.

### Statistical analysis

Statistical significance was determined using Student’s t-test (p<0.05). Error bars represent standard deviation. Minimum of 3 independent biological replicates were conducted for each experiment. Data was analyzed using SPSS Version 21.0 (IBM Corp.).

## Results

### PARP2 expression is serum responsive but is not regulated by its putative SRE

PARP2 is ubiquitously expressed in mammalian cells and a putative serum response element (SRE) within the promoter of the *PARP2* gene suggested its expression is subject to serum stimulation (Ame et al. 2001; Ame et al. 1999). To test whether this SRE was functional, we first assayed PARP protein levels in HeLa, COS-7, MCF-7, NIH3T3 and inner medullary collecting duct (IMCD) cells cultured in complete medium (CM) or serum-free (SF) medium. Immunoblot analysis with the PARP H-250 antibody revealed a prominent 116kDa band corresponding to PARP1 in all cell lines with the exception of NIH3T3 cells (Fig. 1A). A band at 89kDa, corresponding to the C-terminal fragment of PARP1 (Chaitanya et al. 2010) was detected in COS-7 cells and weakly in MCF-7 cells. The appearance of these bands did not change significantly when cells were cultured in CM or SF medium. The H-250 antibody also recognized a 62kDa band, corresponding to PARP2, in all cells cultured in CM (Fig. 1A). This band, however, was absent when COS-7 or MCF-7 cells were cultured in SF medium (Fig. 1A). ERK staining, used as a loading control in this and other studies (Fernandez-Garcia et al. 2007; Rygiel et al. 2008; Xu et al. 2012) showed that protein was present in all samples, with approximately equal amounts assayed in the COS-7 and MCF-7 lanes under the different culturing conditions (Fig. 1).

**Fig. 1.**
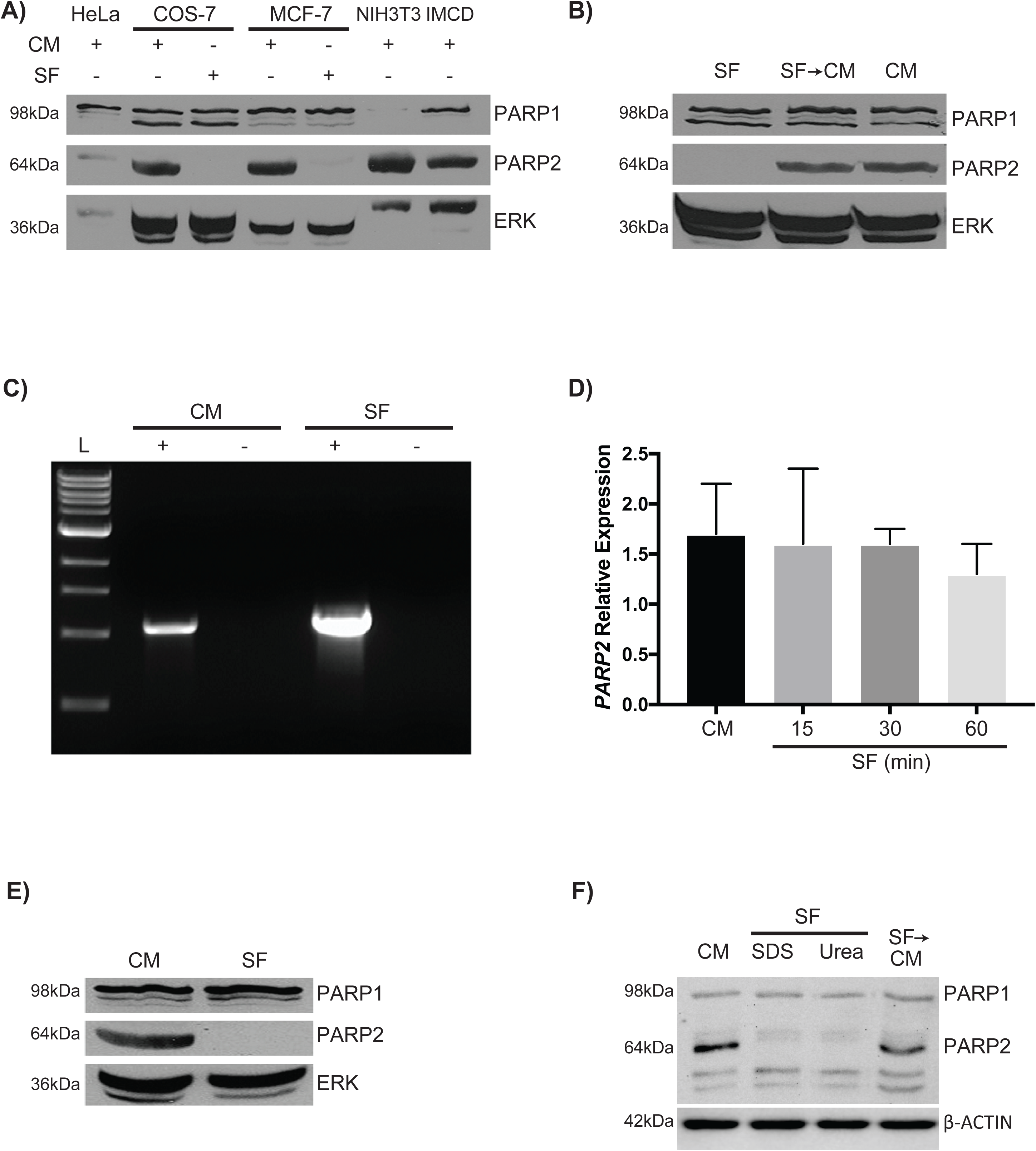
PARP2 is not detected in serum-deprived cells. (A) Immunoblots of protein lysates from HeLa, COS-7, MCF-7, NIH3T3, and IMCD cells cultured in CM and probed with the H-250 PARP1/2 antibody to detect PARP1 (top panel) and PARP2 (middle panel), and with an ERK antibody for a loading control. For the COS-7 and MCF-7 lines, cells were cultured in CM until 90% confluent, and then the medium was replaced with SF medium for 1 hour. (B) The loss of the PARP2 signal in COS-7 cells cultured in SF medium was not due to cell death since adding medium containing serum (CM) (lane 2), allowed cells to reacquire the PARP2 signal. MCF-7 cells were cultured in CM until 90% confluent and then cultured for 30 minutes in either SF medium or maintained in CM. RNA was extracted from each sample and (C) endpoint and (D) quantitative RT-PCR was done with human *PARP2* specific primers. The + lane had reverse transcriptase added to the first strand reaction, and the - lane was the control for genomic contamination. Analysis from qRT-PCR data did not reveal any significant differences between cells cultured in CM compared with those cultured for various times in the absence of serum. (E) Proteins collected from MCF-7 cells cultured as in (C), were used for immunoblotting analysis with antibodies against PARP1/2 and ERK. The reappearance of the PARP2 signal when serum was replaced was indicative that the cells had not undergone apoptosis. (F) Proteins collected from NIH3T3 cells cultured as in (C), were used for immunoblotting analysis with antibodies against PARP1/2 and β-ACTIN. PARP2 signal was absent in SF cells when SDS or urea extractions were used. Unless stated otherwise, results are representative of three independent experiments.

The loss of the PARP2 signal in cells cultured in SF medium was not due to cell death as within one hour, when cultured in CM, these previously serum-starved COS-7 cells had reacquired a PARP2 signal (Fig. 1B). Furthermore, signals were comparable to those in cells that had been continually growing in CM. These results showing that PARP2 levels were influenced by the presence or absence of serum implied that regulation was either at the level of the gene, the protein or both. To address whether or not serum had an effect on altering the activity of the *PARP2* gene, and specifically the SRE in its promoter, MCF-7 cells were cultured in CM or SF medium for 30 minutes and then total RNA was extracted and used for cDNA synthesis. Endpoint and quantitative RT-PCR using *PARP2* primers showed that *PARP2* mRNA was available regardless of the treatment (Fig. 1C, D). However, immunoblot analysis showed PARP2 signals in cells cultured in CM, but not in those that were cultured in SF medium (Fig. 1E).

To address whether the loss of PARP2 signal under SF conditions had physiological effects on cells, we used F9 cells to test whether the loss of PARP2 affected their differentiation potential. F9 cells cultured in CM and treated with RA upregulated the expression of differentiation makers *Gata6* and *Hoxb1* (Fig. S1). Interestingly, F9 cells cultured under SF conditions and treated with RA showed significantly lower expression of *Gata6* and *Hoxb1* when compared to controls (CM + RA), suggesting that loss of PARP2 signal through serum starvation attenuates RA-induced differentiation of F9 cells (Fig. S1).

Together these observations indicating that *PARP2* expression was not affected by serum deprivation brought into question the significance of the putative SRE. More importantly, they strongly suggested that the loss of PARP2 signals could be the consequence of the accelerated degradation of the protein itself or its sequestration to a compartment resistant to Laemmli extraction buffer following serum starvation. To test the latter, 2% SDS and 8M urea lysis buffers were used to lyse NIH3T3 cells cultured under CM or SF conditions. Results show PARP2 signals were absent in both SF cells lysed in 2% SDS and urea lysis buffer (lanes 2 and 3, respectively, Fig. 1F), and indicate that PARP2 is either sequestered in a detergent-insoluble fraction or proteolytically degraded.

### PARP2 is long-lived in cells cultured in complete medium

Since results indicated that PARP2 might be a short-lived protein when cells were cultured in SF medium, an *in silico* analysis was done to identify PEST sequences (mobyle.pasteur.fr), which are responsible for the rapid turnover of many short-lived proteins (Belizario et al. 2008). Although no putative sites were identified in PARP2, COS-7 cells were cultured in CM and protein turnover examined when translation was blocked using cycloheximide (CHX). Cells were cultured in the presence of CHX (50µgml^-1^) for 1, 3, 5 and 7hr, and then processed for immunoblot analysis to detect PARP2 (Fig. 2A). Contrary to the rapid disappearance seen in SF culture, PARP2 appeared stable over the 7hr period. Similar results were seen for PARP1 and ERK, which together would indicate that the loss of the PARP2 signals was serum-dependent.

**Fig. 2.**
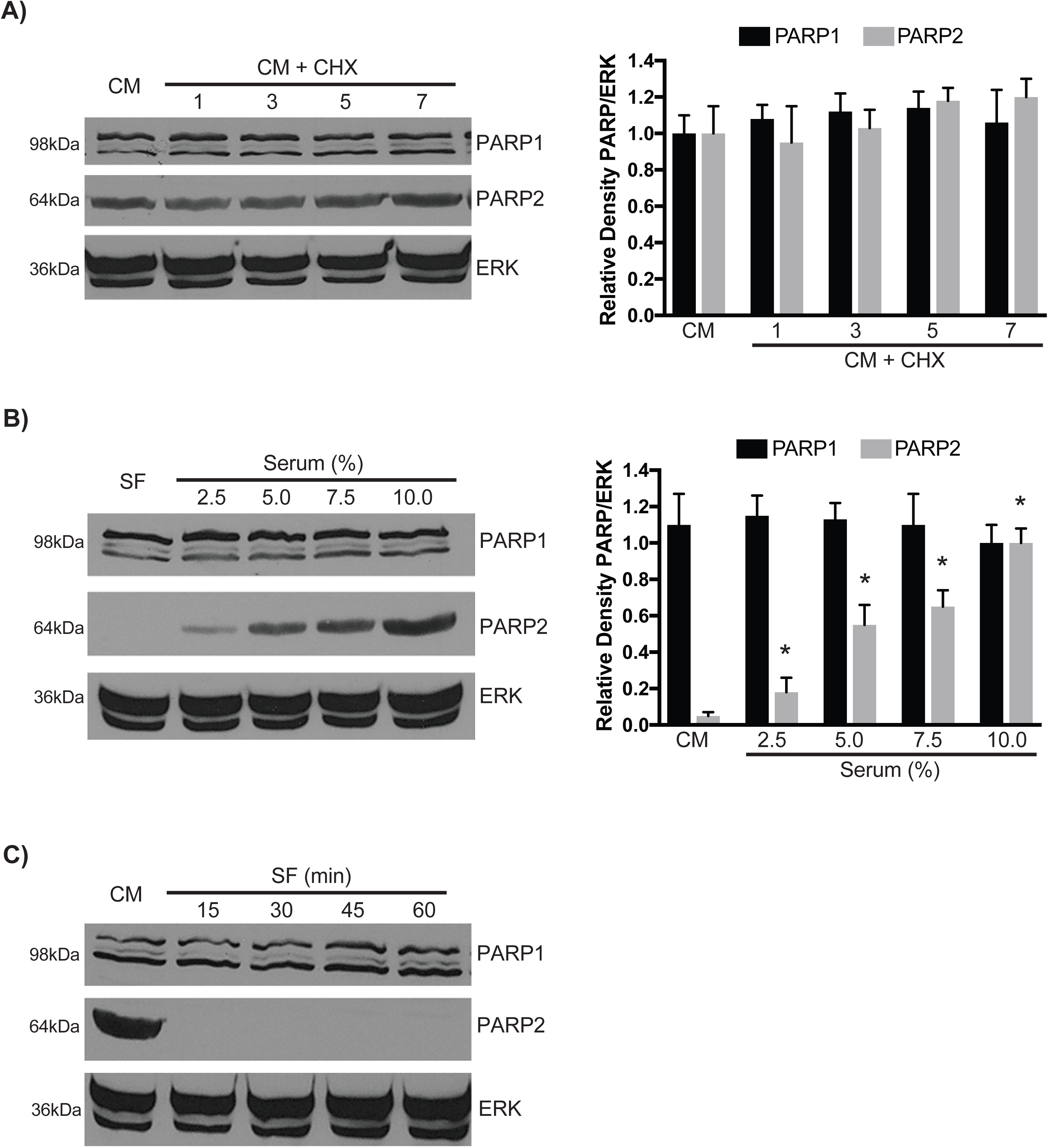
PARP2 is a long-lived serum-dependent protein. (A) COS-7 cells were cultured in CM until 90% confluent, and then cycloheximide (CHX) was added. Cell lysates were collected after 1, 3, 5 or 7hr (lanes 2, 3, 4 and 5, respectively) and probed with antibodies against PARP1/2 and ERK. Results show that there were no significant differences in the PARP1 or PARP2 levels following CHX treatment. (B) COS-7 cells were cultured in SF medium or medium containing increasing amounts of serum as indicated. Immunoblot analysis of lysates probed with antibodies against PARP1/2 and ERK revealed significant increase in levels of PARP2 corresponding to increased serum levels in the medium. (C) COS-7 cells were cultured in CM until 90% confluent, and then the medium was replaced with SF medium. Immunoblot analysis of lysates from cells harvested at 15-minute intervals for 60 minutes and probed with antibodies against PARP1/2 and ERK, show the rapid decline in PARP2 levels following starvation. \**P*<0.05.

The loss of the 62kDa PARP2 signal in cells cultured in SF medium prompted us to undertake a more detailed investigation on the relationship between serum treatment and the levels of PARP2 (Fig. 2B). COS-7 cells were cultured in SF medium or in medium containing increasing amounts of serum. Immunoblot analysis showed a similar staining pattern for full-length PARP1 and its C-terminal fragment, regardless of whether serum was present or not (Fig. 2B). In contrast, increasing the serum concentration from 0 to 10% resulted in the significant increase in the appearance of PARP2 (*P*<0.05; Fig. 2B). Together these results showed that PARP2, but not PARP1, changes in cells in response to serum. Furthermore, the evidence with that seen in figures 1C and 1D, would suggest that this increase is at the protein level rather than due to increased gene activity.

Having determined that PARP2 was available as an SDS soluble protein and this was dependent on the presence of serum, the next question was to address how fast the PARP2 signal would decline when cells were deprived of serum. To determine this, COS-7 cells were cultured in CM for 24hr until 90% confluence, and then the medium was removed and replaced with SF medium. Cell lysates were collected at 15, 30, 45 and 60 minutes and then processed for immunoblot analysis with the H-250 PARP antibody. Results showed PARP1 levels were unaffected by the serum conditions, and comparable signals were seen in all lanes (Fig. 2C). In contrast, PARP2 signals were absent within 15 minutes after serum deprivation (Fig. 2C). This finding suggested serum starvation activated an efficient mechanism to reduce the PARP2 signal, which could be either proteolytic degradation or sequestration to an insoluble fraction.

### PARP2 reduction following serum deprivation is not mediated by a known protease

Since caspase-8 is known to cleave PARP2 in apoptotic murine neurons (Benchoua et al. 2002), and caspase activation is seen in osteoblastic cells following serum deprivation (Mogi et al. 2004), this group of proteases was the first to be investigated. COS-7 cells were treated with 40µM of either the caspase-8-specific inhibitor, Z-Ile-Glu(OMe)-Thr-Asp(OMe)-FMK (IETD) or the broad-spectrum caspase inhibitor Boc-Asp(OMe)-FMK (CI III), and whole cell lysates were collected for immunoblot analysis to detect PARP2 (Fig. 3A). PARP1 levels in cells cultured in SF medium and treated with either of the two inhibitors were comparable to those in cells cultured in CM. The PARP2 signal, however, declined despite the presence of the caspase inhibitors (Fig. 3A). These results, together with those seen in figure 1A, led us to dismiss the notion that the decline in PARP2 levels in cells growing in SF medium was a caspase-dependent, apoptotic-related event.

**Fig. 3.**
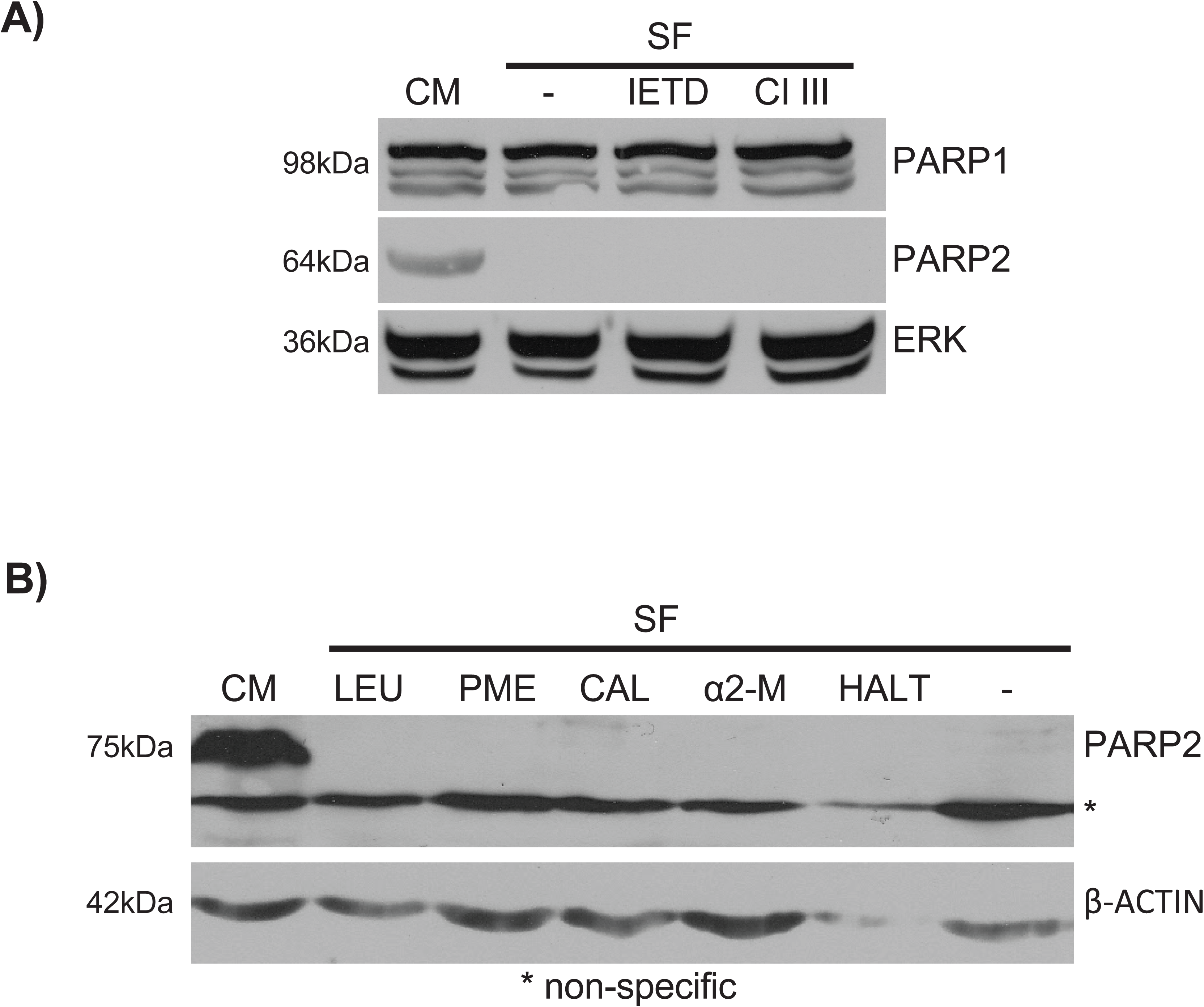
Proteolytic inhibitors do not affect the loss of PARP2 protein. A) COS-7 cells were cultured in CM until 90% confluent and then treated for 5hr with: DMSO vehicle control (lanes 1 and 2), the caspase-8 inhibitor, IETD (lane 3), or the broad-spectrum caspase inhibitor, CI III (lane 4). Following treatment, the medium was replaced with DMSO in CM (lane 1) or SF medium (lane 2), or SF medium containing IETD (lane 3) or CI III (lane 4). Cells were cultured for 15 minutes and then lysates collected for immunoblot analysis with antibodies against PARP1/2 and ERK. Results show that the PARP2 signal disappears when caspase enzymes-inhibited cells were cultured in SF medium. (B) Mouse embryonic fibroblast F20 cells were cultured in CM until 90% confluent, and then for 5hr in CM containing leupeptin (LEU, 250µm), Pepstatin A methyl ester (PME, 30µm), calpeptin (CAL, 30µm), α2-macroglobulin (α2-M, 50µgml^-1^) or 1X HALT protease inhibitor cocktail containing 1.25µM EDTA. Following treatment, the medium was replaced with SF medium containing the same protease inhibitors and the cells cultured for an additional 15 minutes. Cells continually cultured in CM served as controls. Cell lysates from all treatments were collected for immunoblot analysis with an antibody specific to PARP2 (Yucatan) or β-ACTIN. Results show that inhibition of a broad spectrum of proteases was not able to preserve the PARP2 signal in cells cultured in SF medium. The asterisk denotes non-specific staining.

Since the caspase inhibitors had no effect on preventing the disappearance of the PARP2 signal when cells were deprived of serum, the next step was to broaden the search for other proteases that might be involved in the process. MEF F20 (PARP1^+/+^) cells were selected for these studies since the Yucatan PARP2 antibody provided little to no consistent signal with human PARP2 but is robust in detecting mouse PARP2 (data not shown). Furthermore, comparable levels of PARP2 were observed between F20, COS-7 and MCF-7 cells (data not shown). F20 cells were cultured in CM until 90% confluence and then pretreated with 250μM leupeptin (LEU), 30μM pepstatin A methyl ester (PME), 30μM calpeptin (CAL), or 50μgml^-1^ α-2-macroglobulin (α2-M). After 5hr, the medium was replaced with SF medium containing the corresponding protease inhibitor. To inactivate a broad spectrum of endo- and exopeptidases, cells were treated with the 1x HALT Protease Inhibitor Cocktail, which contains AEBSF-HCl, aprotinin, bestatin, E-64, leupeptin, pepstatin A and EDTA. After a 15-minute incubation, cell lysates were collected and processed for immunoblot analysis with the Yucatan PARP2 antibody. Results showed that the inhibitors, either alone or in a cocktail (HALT), were not effective in preventing the disappearance of the PARP2 signal (Fig. 3B). This led us to conclude that under serum deprivation, PARP2 levels were not affected by an amino, serine, cysteine, metallo- and aspartic acid protease. After exhausting a broad spectrum of candidate proteases thought to be responsible for degrading PARP2, our attention turned to the ubiquitin-proteasome system (UPS).

### PARP2 is ubiquitinated

Without finding an inhibitor that would ensure the PARP2 signal was present under SF conditions, and since the UPS is involved in degrading PARP1 (Masdehors et al. 2000; Wang et al. 2008), we considered the same might be true for PARP2. To address this, confocal microscopy was used to examine if EGFP-PARP2 and mRFP-Ubiquitin encoded by vectors transfected into HeLa cells would co-localize. Results showed that PARP2 was present in the nucleus, while ubiquitin was present in the nucleus and cytoplasm (Fig. 4A). Although the co-localization of these proteins in the nucleus was only suggestive that PARP2 was ubiquitinated, further analysis was necessary to provide conclusive evidence. An *in vitro* assay using human recombinant GST-PARP2 and an immunoblot analysis with a GST antibody showed a smear of higher molecular weight PARP2 when Protein Fraction II containing ubiquitination enzymes was present (lane 2, Fig. 4B). Omitting the ubiquitination enzymes served as a negative control (lane 1, Fig. 4B), indicating that PARP2 could be ubiquitinated *in vitro*. To confirm the ubiquitination of PARP2, COS-7 cells were co-transfected with *pBCGST-PARP2* and *pMT123HA-ubiquitin* plasmids, and then cultured in CM. Cells transfected without *pMT123HA-ubiquitin* served as a negative control. Cell lysates were collected, and GST pull down assays were performed prior to immunoblot analysis with an anti-HA antibody. Results showed that ectopically expressed mouse PARP2 was ubiquitinated (lane 1, Fig. 4C), and the amount of ubiquitin (smear) accumulated following MG-132 treatment (lane 2, Fig. 4C). As expected, no HA-ubiquitin signal was detected in cells transfected with *pBCGST-PARP2* alone (lane 3, Fig. 4C). A Simian virus 40 nuclear localization signal epitope tagged to GST served as a positive control and was also ubiquitinated (lane 4, Fig. 4C). Thus, when serum was present, exogenously expressed PARP2 was ubiquitinated and despite being a long-lived protein under these conditions (Fig. 2A), this post-translational modification targets PARP2 to the proteasome.

**Fig. 4.**
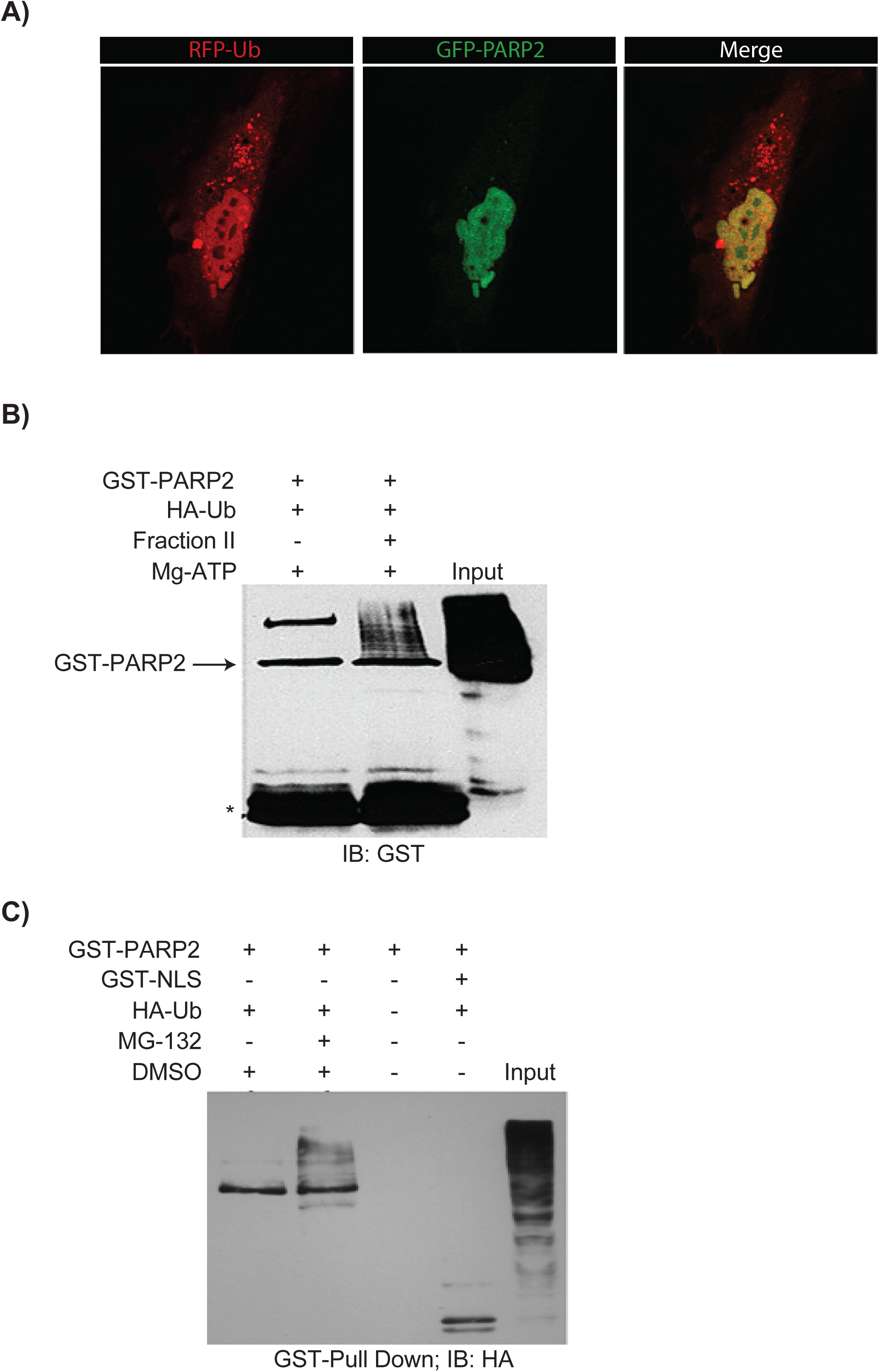
PARP2 protein is ubiquitinated. (A) HeLa cells were transfected with *pEGFP-PARP2* and *RFP-ubiquitin* and then cultured in CM for 24hr. To visualize the fluorescence, cells were fixed in 4% paraformaldehyde in PBS, mounted in ProLong Gold antifade medium, and viewed on a Zeiss LSM 510 confocal microscope. Results show PARP2 staining in the nucleus (panel B). Ectopically expressed ubiquitin was seen in the cytoplasm and nucleus (panel A), and the overlap in PARP2 and ubiquitin staining is seen in yellow when the images were merged (panel C). (B) Human GST-PARP2 recombinant protein/Mg-ATP +/- Ubiquitin was incubated without Fraction II (lane 1) or with Fraction II (lane 2) and resolved by SDS-PAGE. Proteins were transferred to blots and probed with a GST-specific antibody. The smear of higher molecular weight PARP2 seen in lane 2 was indicative that the protein had been ubiquitinated by enzymes in Fraction II. The band around 150kD in lane 1 was GST-PARP2 dimers present in commercial recombinant GST-PARP2 protein. The intense signal seen in the input lane was GST-PARP2 smear from COS-7 cells overexpressing *pBC-GST-PARP2*. (C) COS-7 cells were transfected with *pBCGST-PARP2* or *pBC-GST-NLS*, with or without *pMT123HA-ubiquitin* and then treated with MG-132 or left untreated as a control. Cells were lysed in 1x RIPA buffer and the lysates used in GST-pull down assays. Results show that PARP2 was ubiquitinated (lane 1) and the levels accumulated when proteasome activity was inhibited with MG-132 (lane 2). The input lane was protein from COS-7 cells that were overexpressing GST-PARP2 and HA-ubiquitin. Cells transfected with *pBC-GST-PARP2* alone did not show an HA signal (lane 3), while the Simian virus 40 nuclear localization signal and positive control was ubiquitinated (lane 4). The GST immunoblot (right panel) confirmed the presence of the GST-epitope tagged proteins in the samples probed with the HA antibody (left panel). The asterisk represents GST fragments from the recombinant proteins.

### PARP2 is degraded by the proteasome in the presence of serum

The ubiquitination results, together with those showing that the proteasome inhibitor MG-132 prevented ectopically expressed PARP2 from degrading, strongly suggested that the UPS was the mechanism used to regulate endogenous PARP2 levels. Unexpectedly, however, results showed that MG-132 had no effect on preventing endogenous PARP2 from being detected under SDS-denaturing conditions in cells cultured under SF conditions (Fig. 5A). As a comparison, immunoblot analysis showed COS-7 cells cultured in CM and treated with MG-132 had consistent PARP2 signals (Fig. 5B). In fact, the signals increased with increasing concentrations of MG-132, which was confirmed by densitometric analysis showing the significant increase over levels in cells cultured in CM alone. Although PARP1 was reported to be ubiquitinated and degraded in the proteasome (Ame et al. 2009), we did not see any appreciable change after MG-132 treatment, which mimicked that seen under SF conditions. Together this data would indicate that when cells were cultured in CM, endogenous PARP2 is ubiquitinated and degraded in the proteasome. The data also points to the fact that different mechanisms of PARP2 regulation exist and that these processes are activated in a manner that depends on the conditions under which the cells are cultured.

**Fig. 5.**
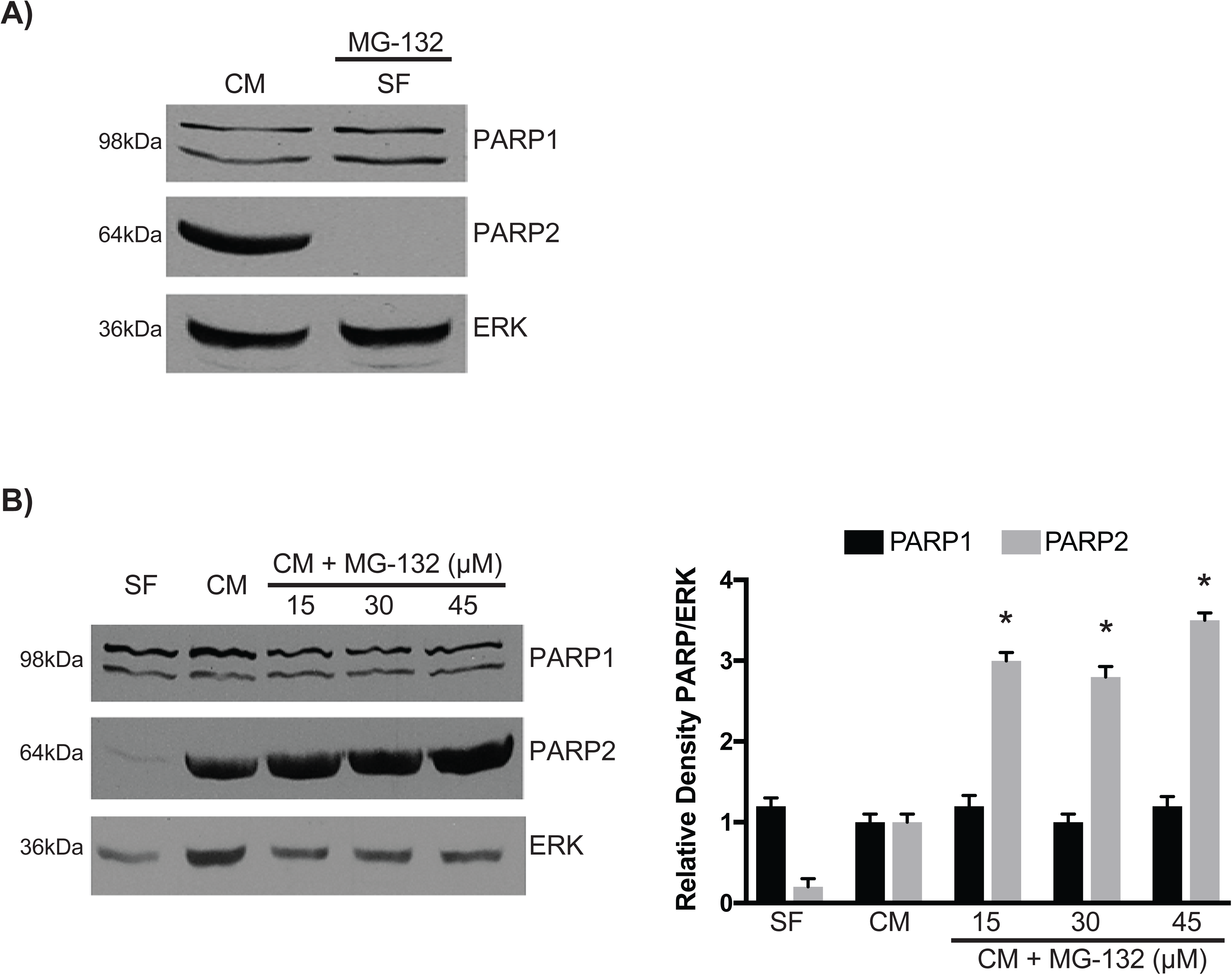
The ubiquitin-proteasome system (UPS) is involved in PARP2 degradation under serum conditions. (A) COS-7 cells were cultured in CM until 90% confluent and then transferred and cultured for 5hr in CM containing MG-132. Cells were cultured for 15 minutes in SF medium containing MG-132, and lysates collected for immunoblot analysis with antibodies against PARP1/2 and ERK. Inactivating proteasome activity had no effect on preserving the PARP2 signal. (B) COS-7 cells were cultured in CM for 24hr and then in SF (lane 1), CM (lane 2), or CM containing different concentrations of MG-132 (lanes 3-5). Immunoblot analysis with antibodies against PARP1/2 and ERK of lysates from cells maintained for 24hr under these conditions shows significant increase in levels of PARP2, but only in cells cultured in CM with MG132. \**P*<0.05.

### PARP2 is sequestered to an insoluble fraction after serum deprivation

Having shown PARP2 is degraded through the UPS in cells cultured in CM and having excluded the involvement of many known protease types responsible for its absence after serum withdrawal, we set out to examine if PARP2 was sequestered to an insoluble compartment following serum deprivation. If PARP2 was degraded rather than being sequestered after serum withdrawal, then it was expected that the signals would not reappear, as indicated in figure 1B, after serum was added and protein synthesis inhibited with CHX.

Immunoblot analysis with a PARP2 antibody (Axxora) showed, as expected, the 62kDA PARP2 signal in cells cultured in CM (Fig. 6A, lane 1). The PARP2 signal was lost when serum was removed and remained so in cells cultured in SF medium and treated with CHX (Fig. 6A, lanes 2 and 3, respectively). Similarly, CHX had no apparent effect on PARP2 when cells were cultured in CM (Fig. 6A, lane 4). However, when cells were treated with CHX, and cultured under serum-free conditions, were transferred to CM containing CHX, the PARP2 signal appeared (Fig. 6A, lane 5). In fact, the signal strength was almost identical to cells cultured without CHX (Fig. 6A, lane 6). These results would suggest serum starvation led to the sequestration of PARP2 to an SDS- (and urea) insoluble compartment, which could recycle back to a soluble form when serum was present. To further understand the capacity of this sequestration, we overexpressed GST-PARP2 in MCF-7 cells cultured in CM or SF medium for 15, 30 and 60 minutes. Immunoblot analysis of these lysates, using a GST antibody, showed that the GST-PARP2 signal was comparable between the cells cultured in CM and SF medium (arrow, Fig. 6B). Lower molecular weight signals were also seen on blots (arrowheads, Fig. 6B), and although not characterized, they may be PARP2 fragments that appeared due to ubiquitin-mediated proteasomal degradation in CM. Thus, the evidence would suggest that either the sequestration capacity was limiting, or the GST tag conferred solubility to PARP2, either way with most of the ectopically expressed PARP2 free in the SDS-soluble fraction. Together, the results indicate that serum withdrawal does not lead to PARP2 degradation.

**Fig. 6.**
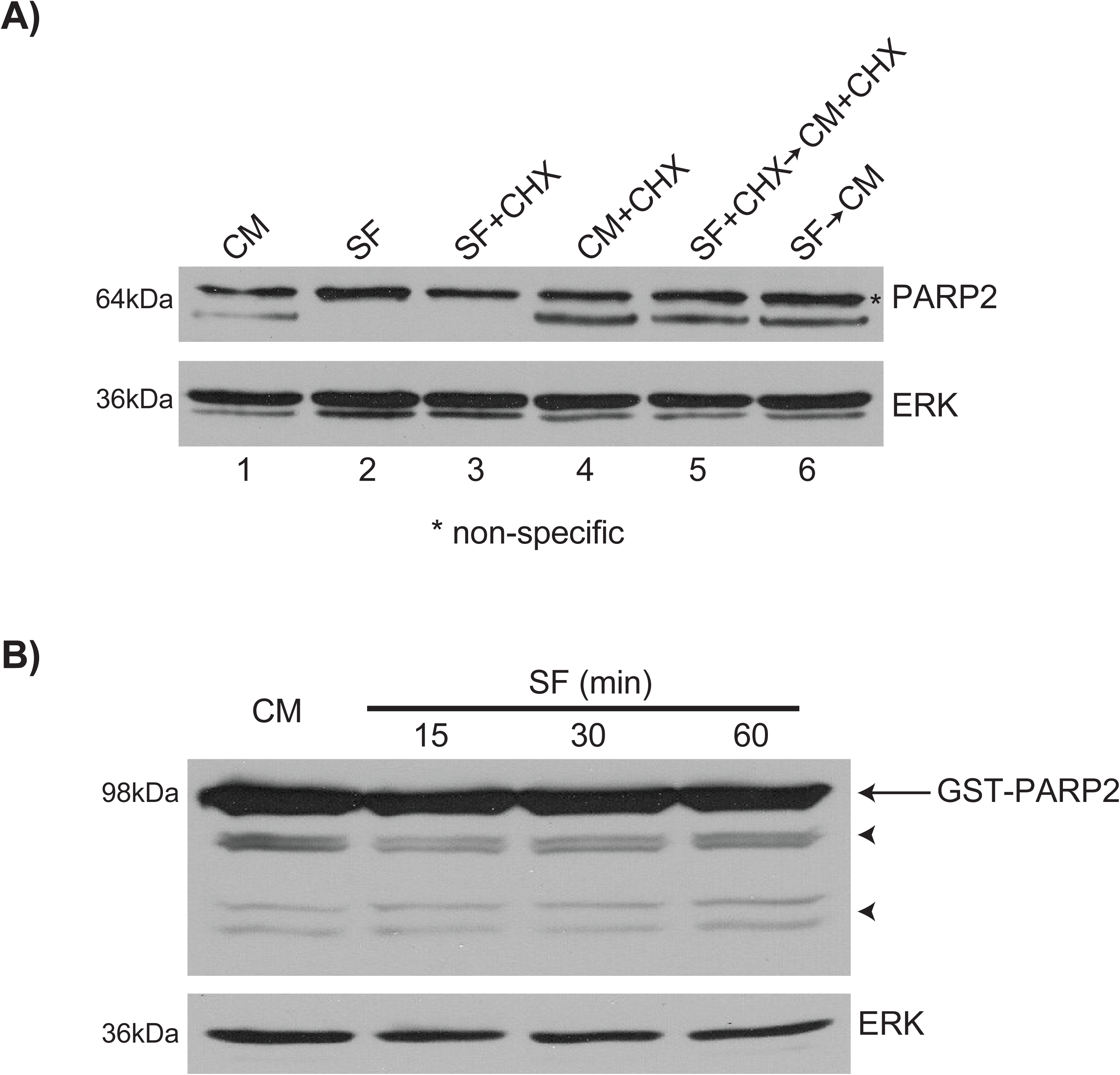
PARP2 is sequestered to a SDS-insoluble fraction following serum starvation. (A) MCF-7 cells were cultured in CM until 90% confluent and then medium was replaced with SF medium containing CHX or in CM with CHX. After 7hr, cells in lanes 1-4 were lysed in 1x Laemmli extraction buffer. Cells in lanes 5 and 6 were cultured in CM or CM and CHX, respectively, for an additional hour and then lysed in Laemmli buffer. All samples were probed with antibodies against PARP2 (Axxora) and ERK. Though no apparent differences were seen in the PARP2 signals between samples in lanes 4-6, the presence of a band in lane 5, under conditions where protein synthesis was inhibited, would indicate that PARP2 had been sequestered into an SDS-insoluble fraction resulting from serum deprivation. (B) MCF-7 cells were transfected with *pBC-GST-PARP2* and after 24hr, CM was replaced with SF medium. Cells were serum starved for 15, 30 or 60 minutes, and analyzed with antibodies against GST and ERK, show that the GST-PARP2 signal was comparable between the cells cultured in CM and SF medium (arrow). The asterisk in panel A denotes non-specific staining, and minor bands seen on blots (arrowheads, panel B) may be PARP2 fragments from ubiquitin-mediated proteasomal degradation in CM.

## Discussion

PARP proteins were first identified as being players in DNA repair (Gupte et al. 2017; Wei and Yu 2016), and this involves initiating their poly ADP-ribosylation polymerase activity (Barzilai and Yamamoto 2004; Dantzer et al. 2006a; Huber et al. 2004). In addition to this crucial role, PARP proteins serve other functions involved in, but not limited to the regulation of gene activity, cell death, the immune system, cellular metabolism and differentiation (Bai 2015; Vida et al. 2017). PARP2 is required for initiating the differentiation of F9 cells into primitive extraembryonic endoderm (Quenet et al. 2008), and our results showing *Gata6* and *Hoxb1* expression under serum-free conditions (Fig. S1) would indicate that PARP1 was unable to act in a functionally redundant manner (Quenet et al. 2008) and implicating PARP2 in differentiation.

Whatever the case, and given this diversity, the mechanism(s) to control the level and activity of these enzymes must be tightly regulated. In the case of PARP1 and PARP2, basal activities are low, while in response to DNA damage this changes rapidly (Bai and Canto 2012; Krishnakumar and Kraus 2010; Langelier et al. 2014). At the level of the gene, the presence of a putative serum response element (SRE) in the *PARP2* promoter region (Ame et al. 2001) suggested that transcriptional activity might be influenced by growth factor and/or mitogen stimulation. To investigate this, we cultured different cell lines in the presence or absence of serum and then assayed PARP2 levels. The loss of the PARP2 signal in cells cultured in SF medium (Fig. 1A), and its return when serum was replaced (Fig. 1B) supported the notion that the *PARP2* gene was serum responsive. Given this on-off appearance at the protein level, and the presence of the putative SRE, we had expected to see changes in *PARP2* mRNA expression under the different culturing conditions. This, however, was not the case and the presence of a *PARP2* amplicon in cells cultured in the SF medium indicated that the message was available (Fig. 1C). Furthermore, the fact that the quantitative-RT-PCR results showed no significant differences in expression in cells cultured under the different conditions (Fig. 1D) indicated that the regulation of PARP2 in cells deprived of serum was not due to the direct regulation of the gene, at least within the time frame of our investigation. Thus, the presence of *PARP2* mRNA under the SF conditions did not seem to contribute to the fast return (within one hour) of the PARP2 signal after serum was added to cells incubated in CHX to block new protein synthesis (Fig. 6A). Also, the reappearance of the PARP2 signal when serum was added was dose-dependent (Fig. 2B) and not reliant on increased transcription that was dependent on the putative SRE in the *PARP2* promoter (Fig. 1D). These results implied that PARP2 under serum-free conditions was either being rapidly degraded (Fig. 2C) or sequestered in an insoluble compartment.

Serum withdrawal in different cultured cells activates many proteolytic, and proteolytic-related proteins including caspases, calpains, autophagy-related proteins, ubiquitin, and proteasome subunits (Fuertes et al. 2003; Kilic et al. 2002; Mogi et al. 2004; Nakashima et al. 2005; Schamberger et al. 2005). The fact that caspases can cleave PARP2 (Benchoua et al. 2002) placed this family of cysteine-dependent aspartate-directed proteases at the forefront of candidates responsible for the serum-dependent changes seen with the different cell types (Fig. 1). Unfortunately, our analysis using a caspase-specific and a broad-spectrum caspase inhibitor (Fig. 3A) as well as a proteolytic inhibitor cocktail (Fig. 3B) ruled out the possibility that caspases were responsible for the proteolysis. Subsequent experiments were designed to explore the ability of serine, cysteine, metallo- and aspartic proteases to alter PARP2 levels (Fig. 3B). As with the caspase inhibitors, those routinely employed to prevent proteolysis did not have an effect on the PARP2 signals when cells were cultured in serum-free conditions (Fig. 3B). Furthermore, the reports that the cysteine protease cathepsin L is present in the nucleus, like PARP2, and is involved in cell-cycle progression (Puchi et al. 2010) prompted us to block its activity to see what effect it would have on endogenous PARP2 levels in serum-deprived cells. Leupeptin, an inhibitor of endosomal trypsin-like serine and cysteine proteases (Simmons et al. 2005) had no effect, which ruled out the involvement of cathepsin L. Details of the inhibitor studies suggested that the disappearance of the PARP2 signal may not be the result of proteolytic degradation, but instead and as noted above, due to the sequestration of the protein to an insoluble form under serum-free conditions (Fig. 2B).

The detergent solubility of constituents within a cell is determined by the chemical properties of the substrates as well as the detergents. Some cellular entities are naturally resistant to extraction by detergents (Horigome et al. 2008; Takata et al. 2009) while other proteins may be converted to detergent-soluble or insoluble forms following various stimulations (Peters et al. 2012; Reis-Rodrigues et al. 2012). For instance, serum starvation causes the translocation of dynein to a more detergent-soluble compartment in NRK cells, and this change is reversed when serum is added (Lin et al. 1994). Serum withdrawal also leads to sequestration of caspase-9 into detergent-insoluble cytoskeletal structures in rat 423-cells (Schamberger et al. 2005). The same is true for IM-9 cells, where detergent insolubility of growth hormone receptors occurs due to ligand-induced formation of cross-linked disulfide bonds (Goldsmith et al. 1997). These reports and our results led us to investigate if serum deprivation caused an SDS solubility change in PARP2. Furthermore, they raised the question of what in serum ameliorates the SDS solubility of PARP2? Conversely, why does PARP2 become SDS insoluble when cells are serum-deprived and what process initiates this sequestration or biochemical change? More importantly, what physiological function does this regulation serve in cells, or does it even have biological significance? Despite the similarities between PARP1 and PARP2, that the solubility in SDS of the former did not change following serum deprivation would indicate that serum withdrawal activated a PARP2-specific mechanism.

PARP1 regulation has been studied extensively and the protein can be found in lamin-enriched or DNA-bound detergent-resistant fractions (Frouin et al. 2003; Vidakovic et al. 2004). Furthermore, PARP1 changes its solubility in NP-40 detergent when modified by sumoylation following heat shock (Martin et al. 2009). Several other proteins involved in regulating DNA replication and repair, e.g. PCNA, P21, OGG1, XRCC1 and CAF-1 P150 are also found in DNA-bound detergent-resistant fractions (Amouroux et al. 2010; Campalans et al. 2013; Frouin et al. 2003; Okano et al. 2003). Although PARP2 functions in DNA repair, this association with DNA would only enhance its resistance to some nonionic detergents including Triton and NP-40, but not to anionic detergents like SDS. Thus, despite the link between serum starvation to activate the DNA damage response pathway in some cancer cells and to induce DNA fragmentation in normal cells (Lu et al. 2008; Shi et al. 2012), and based on the chemistry noted above, it is questionable that PARP2 would be insoluble to SDS (and urea) if serum starvation had caused it to bind to DNA lesions.

PARP2 can also localize to the cytosol in gonocytes, spermatogonia and spermatids in mice (Gungor-Ordueri et al. 2014), and proteins are known to change their detergent solubility when associated with either glycosylphosphosphatidyl inositol enriched microdomains, the cytoskeleton or when posttranslationally modified (Brown and Rose 1992; Fujita et al. 2011; Ledesma et al. 1994; Paladino et al. 2002; Refolo et al. 1991; Waelter et al. 2001). Moreover, under pathological conditions proteins can become SDS-insoluble, as in the mouse model of Alzheimer’s disease where amyloid β protein (Aβ) changes to SDS-insoluble forms of Aβ42 and Aβ40 (Kawarabayashi et al. 2001) or in Huntington’s disease where the mutated Huntingtin protein with its polyglutamine repeat expansion is resistant to SDS extraction (Heiser et al. 2000; Scherzinger et al. 1999). These conversions to SDS-insoluble forms are disease-dependent and the combinatory effects of conformational changes and oligomerization and fibril formation that appear irreversible in patients with specific neurodegenerative diseases (Cruz et al. 1997; Diaz-Hernandez et al. 2005; Dolev and Michaelson 2004; Wong et al. 2008). The conversion of PARP2 between soluble and insoluble forms under different serum conditions that we observed suggests the protein adopts a physiological conformation in either condition, linked to a stress response. This is evident in nematodes and yeast where the accumulation of SDS-insoluble proteins in cells is indicative of aging (Peters et al. 2012; Reis-Rodrigues et al. 2012). The accumulation of SDS-insoluble proteins is accelerated by nitrogen starvation even in young yeast cells, where Tor1 kinase plays a regulatory role in this accumulation of a novel autophagic cargo preparation process (Peters et al. 2012). It is unlikely, however, that the loss or sequestration of PARP2 is a mechanism to prepare it for autophagic degradation since inhibiting lysosomal enzymes with leupeptin had no apparent effect (Fig. 3B). Serum deprivation and the way cells cope, however, is indicative of a stress response, and although our results suggest that PARP2 participates in these events, it does not explain why the protein cannot be detected following urea denaturation (Fig. 1F). Urea is often used to recover proteins from inclusion bodies (Burgess 2009), but there is no single method of solubilization for every protein (Singh et al. 2015).

Given the shortcomings on being unable to find a sequestration mechanism noted above, our data does indicate that PARP2 is degraded through the UPS in cells when serum is present (Fig. 5B). PARP1, the structural and functional relative of PARP2 is ubiquitinated and degraded through the 26S proteasome, and the ubiquitination site is mapped to its N-terminal DNA-binding domain (Wang et al. 2008). Two E3 ubiquitin ligases, Iduna and CHFR, ubiquitinate and target PARP1 for proteasomal degradation in a poly-ADP-ribose (PAR)-dependent manner (Kang et al. 2011; Kashima et al. 2012; Liu et al. 2013). Interestingly, Iduna binds and ubiquitinates a panel of DNA damage repair proteins including PARP2 (Kang et al. 2011). It is not known, however, if CHFR ubiquitinates PARP2, but it is recruited to DNA lesions through binding to the PAR moiety of pARylated PARP1, and is the first E3 ligase engaged in protein ubiquitination at DNA damage sites (Liu et al. 2013). When this occurs PARP1 dissociates from DNA lesions and is subsequently degraded in the proteasome. This degradation prevents cells from ATP deprivation due to persistent poly-ADP-ribosylation (pARylation) (Liu et al. 2013) that occurs even by ubiquitinated PARP1 (Wang et al. 2008). Although PARP2 accounts for only 10-15% of the total PARP activity in cells (Bai and Canto 2012), the remaining pARylation and recruitment of CHFR to DNA lesions seen in PARP1 knockdown cells (Liu et al. 2013) is likely the result of other DNA-dependent PARPs, possibly PARP2. If so and in response to DNA damage, the presence of autoPARylated PARP2 may be the result of it binding through the PBZ domain in CHFR. As attractive as this sounds, as serum removal would have initiated mechanisms for cell cycle arrest (Bertoli et al. 2013), it is unlikely with our cell lines that PARP2 ubiquitination and degradation contributes to cell cycle arrest, as noted for CHFR-mediated PARP1 (Kashima et al. 2012). Our rationale for disputing the degradation of PARP2 under these conditions comes from the MG-132 studies (Fig. 5A), where cells should have retained some of the PARP2 protein when they were transferred to serum-free medium.

In summary, PARP2 was shown to be a long-lived protein that is continually degraded by the ubiquitin proteasome system in cells cultured in medium containing serum. However, in the case of a cell stress response when serum is removed, PARP2, through an unknown mechanism, is no longer in a SDS or urea soluble form, and it is not known if this is a prelude to apoptosis or to some other physiological requirement. Nevertheless, alleviating this cellular stress by the addition of serum recycles PARP2 to a form, thereby allowing it to resume its enzymatic role involved in the pARylation of target substrates.

## Acknowledgements

This paper was supported by funds from the Natural Sciences and Engineering Research Council of Canada (NSERC) to GMK. MIG acknowledges support from the Faculty of Graduate and Postdoctoral Studies, University of Western Ontario, the Collaborative Graduate Specialization in Developmental Biology, University of Western Ontario, the Child Health Research Institute and NSERC for a CGS D scholarship. We would also like to thank members past and present of the Kelly lab for discussions, and especially Amy R. Assabgui for contributions to the figures, and the following for generously providing reagents used in this study: Dr. V. Schreiber (École Supérieure de Biotechnologie Strasbourg); Dr. D. Bohmann (University of Rochester); and Dr. G. Poirier (Université Laval).

## Supplementary Figure legend

**S1.**
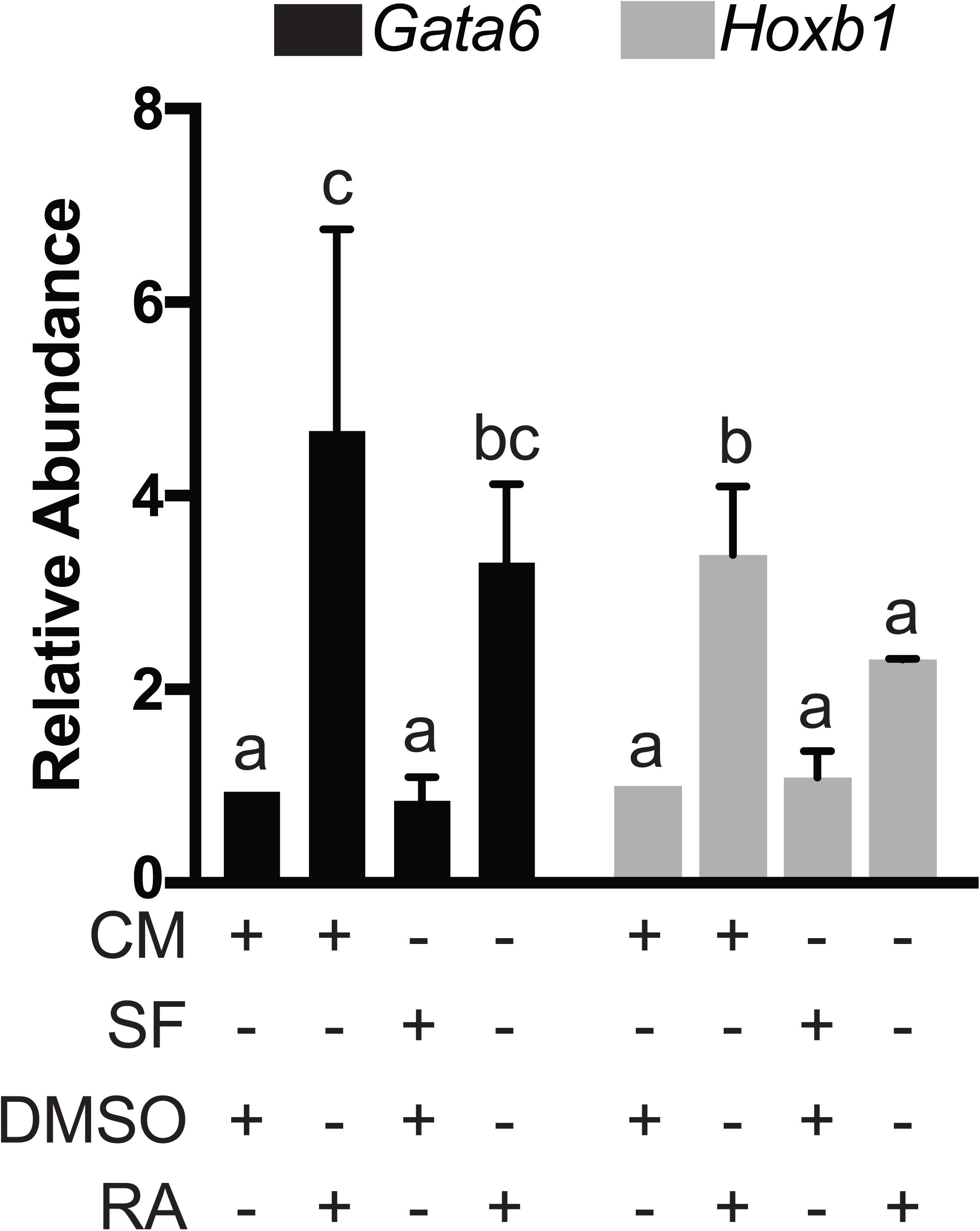
Loss of PARP2 under SF conditions attenuates RA-induced differentiation of F9 cells. F9 cells were cultured in CM for 24 hours and then cultured for 60 minutes in either SF medium or maintained in CM. Media was then changed to either contain DMSO or RA, and either in CM or SF conditions for an additional 12 hours. RNA was extracted from each sample and quantitative RT-PCR was done with mouse *Gata6* and *Hoxb1*-specific primers. Analysis of quantitative RT-PCR showed that F9 cells treated with RA under SF conditions had significantly lower *Gata6* and *Hoxb1* expression.

## Graphical Abstract Legend

**Fig. 1.**
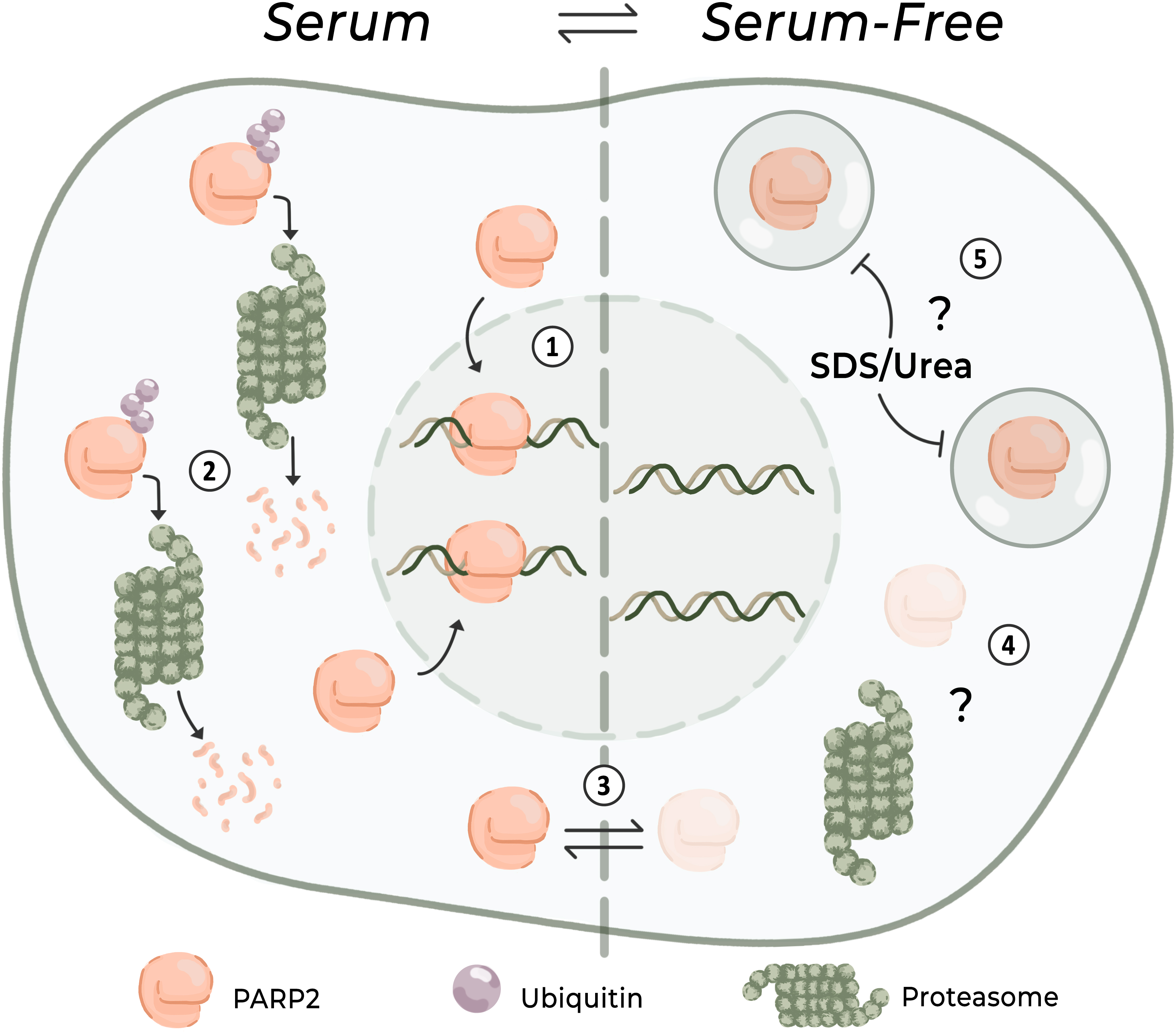
Schematic representation of PARP2 regulation under different growing conditions. PARP2 maintains genomic stability by recruiting DNA repair machinery to sites of DNA damage (1), and when not required is ubiquitinated and degraded by the proteasome (2). Under stress condition (3), such as serum-starvation, PARP2 is either degraded in a proteasome-independent manner (4) or packaged into a SDS/urea-insoluble fraction (5).

## References

Ali, S.O., Khan, F.A., Galindo-Campos, M.A., and Yelamos, J. 2016. Understanding specific functions of PARP-2: new lessons for cancer therapy. Am. J. Cancer Res. 6(9): 1842–1863. Available from https://www.ncbi.nlm.nih.gov/pubmed/27725894.

Ame, J.C., Schreiber, V., Fraulob, V., Dolle, P., de Murcia, G., and Niedergang, C.P. 2001. A bidirectional promoter connects the poly(ADP-ribose) polymerase 2 (PARP-2) gene to the gene for RNase P RNA. structure and expression of the mouse PARP-2 gene. J. Biol. Chem. 276(14): 11092–11099. doi:10.1074/jbc.M007870200.

Ame, J.C., Hakme, A., Quenet, D., Fouquerel, E., Dantzer, F., and Schreiber, V. 2009. Detection of the nuclear poly(ADP-ribose)-metabolizing enzymes and activities in response to DNA damage. Methods Mol. Biol. 464: 267–283. doi:10.1007/978-1-60327-461-6_15.

Ame, J.C., Rolli, V., Schreiber, V., Niedergang, C., Apiou, F., Decker, P., et al. 1999. PARP-2, A novel mammalian DNA damage-dependent poly(ADP-ribose) polymerase. J. Biol. Chem. 274(25): 17860–17868. doi:10.1074/jbc.274.25.17860.

Amouroux, R., Campalans, A., Epe, B., and Radicella, J.P. 2010. Oxidative stress triggers the preferential assembly of base excision repair complexes on open chromatin regions. Nucleic Acids Res. 38(9): 2878–2890. doi:10.1093/nar/gkp1247.

Bai, P. 2015. Biology of Poly(ADP-Ribose) Polymerases: The Factotums of Cell Maintenance. Mol. Cell 58(6): 947–958. doi:10.1016/j.molcel.2015.01.034.

Bai, P., and Canto, C. 2012. The role of PARP-1 and PARP-2 enzymes in metabolic regulation and disease. Cell Metab. 16(3): 290–295. doi:10.1016/j.cmet.2012.06.016.

Bai, P., Houten, S.M., Huber, A., Schreiber, V., Watanabe, M., Kiss, B., et al. 2007. Poly(ADP-ribose) polymerase-2 controls adipocyte differentiation and adipose tissue function through the regulation of the activity of the retinoid X receptor/peroxisome proliferator-activated receptor-gamma heterodimer. The Journal of biological chemistry 282(52): 37738–37746. doi:10.1074/jbc.M701021200.

Bai, P., Canto, C., Brunyanszki, A., Huber, A., Szanto, M., Cen, Y., et al. 2011. PARP-2 regulates SIRT1 expression and whole-body energy expenditure. Cell. Metab. 13(4): 450–460. doi:10.1016/j.cmet.2011.03.013.

Barzilai, A., and Yamamoto, K. 2004. DNA damage responses to oxidative stress. DNA repair 3(8): 1109–1115. doi:10.1016/j.dnarep.2004.03.002.

Belizario, J.E., Alves, J., Garay-Malpartida, M., and Occhiucci, J.M. 2008. Coupling caspase cleavage and proteasomal degradation of proteins carrying PEST motif. Current protein & peptide science 9(3): 210–220. doi:10.2174/138920308784534023.

Benchoua, A., Couriaud, C., Guegan, C., Tartier, L., Couvert, P., Friocourt, G., et al. 2002. Active caspase-8 translocates into the nucleus of apoptotic cells to inactivate poly(ADP-ribose) polymerase-2. J. Biol. Chem. 277(37): 34217–34222. doi:10.1074/jbc.M203941200.

Bertoli, C., Skotheim, J.M., and de Bruin, R.A. 2013. Control of cell cycle transcription during G1 and S phases. Nat. Rev. Mol. Cell. Biol. 14(8): 518–528. doi:10.1038/nrm3629.

Boudra, M.T., Bolin, C., Chiker, S., Fouquin, A., Zaremba, T., Vaslin, L., et al. 2015. PARP-2 depletion results in lower radiation cell survival but cell line-specific differences in poly(ADP-ribose) levels. Cellular and molecular life sciences: CMLS 72(8): 1585–1597. doi:10.1007/s00018-014-1765-2.

Brown, D.A., and Rose, J.K. 1992. Sorting of GPI-anchored proteins to glycolipid-enriched membrane subdomains during transport to the apical cell surface. Cell 68(3): 533–544. doi:10.1016/0092-8674(92)90189-J.

Burgess, R.R. 2009. Refolding solubilized inclusion body proteins. Methods Enzymol. 463: 259–282. doi:10.1016/S0076-6879(09)63017-2.

Campalans, A., Kortulewski, T., Amouroux, R., Menoni, H., Vermeulen, W., and Radicella, J.P. 2013. Distinct spatiotemporal patterns and PARP dependence of XRCC1 recruitment to single-strand break and base excision repair. Nucleic Acids. Res. 41(5): 3115–3129. doi:10.1093/nar/gkt025.

Celik-Ozenci, C., and Tasatargil, A. 2013. Role of poly(ADP-ribose) polymerases in male reproduction. Spermatogenesis 3(2): e24194. doi:10.4161/spmg.24194.

Chaitanya, G.V., Steven, A.J., and Babu, P.P. 2010. PARP-1 cleavage fragments: signatures of cell-death proteases in neurodegeneration. Cell communication and signaling: CCS 8: 31. doi:10.1186/1478-811X-8-31.

Chen, Q., Kassab, M.A., Dantzer, F., and Yu, X. 2018. PARP2 mediates branched poly ADP-ribosylation in response to DNA damage. Nat. Commun. 9(1): 3233. doi:10.1038/s41467-018-05588-5.

Cruz, L., Urbanc, B., Buldyrev, S.V., Christie, R., Gomez-Isla, T., Havlin, S., et al. 1997. Aggregation and disaggregation of senile plaques in Alzheimer disease. Proceedings of the National Academy of Sciences of the United States of America 94(14): 7612–7616. doi:10.1073/pnas.94.14.7612.

Dantzer, F., and Santoro, R. 2013. The expanding role of PARPs in the establishment and maintenance of heterochromatin. FEBS J 280(15): 3508–3518. doi:10.1111/febs.12368.

Dantzer, F., Ame, J.C., Schreiber, V., Nakamura, J., Menissier-de Murcia, J., and de Murcia, G. 2006a. Poly(ADP-ribose) polymerase-1 activation during DNA damage and repair. Methods Enzymol. 409: 493–510. doi:10.1016/S0076-6879(05)09029-4.

Dantzer, F., Giraud-Panis, M.J., Jaco, I., Ame, J.C., Schultz, I., Blasco, M., et al. 2004. Functional interaction between poly(ADP-Ribose) polymerase 2 (PARP-2) and TRF2: PARP activity negatively regulates TRF2. Molecular and cellular biology 24(4): 1595–1607. doi:10.1128/MCB.24.4.1595-1607.2004.

Dantzer, F., Mark, M., Quenet, D., Scherthan, H., Huber, A., Liebe, B., et al. 2006b. Poly(ADP-ribose) polymerase-2 contributes to the fidelity of male meiosis I and spermiogenesis. Proceedings of the National Academy of Sciences of the United States of America 103(40): 14854–14859. doi:10.1073/pnas.0604252103.

Deol, G.S.J., Cuthbert, T.N., Gatie, M.I., Spice, D.M., Hilton, L.R., and Kelly, G.M. 2017. Wnt and hedgehog signaling regulate the differentiation of F9 cells into extraembryonic endoderm. Front. Cell Dev. Biol. 5: 93. doi:10.3389/fcell.2017.00093.

Diaz-Hernandez, M., Torres-Peraza, J., Salvatori-Abarca, A., Moran, M.A., Gomez-Ramos, P., Alberch, J., et al. 2005. Full motor recovery despite striatal neuron loss and formation of irreversible amyloid-like inclusions in a conditional mouse model of Huntington’s disease. J. Neurosci. 25(42): 9773–9781. doi:10.1523/JNEUROSCI.3183-05.2005.

Dolev, I., and Michaelson, D.M. 2004. A nontransgenic mouse model shows inducible amyloid-beta (Abeta) peptide deposition and elucidates the role of apolipoprotein E in the amyloid cascade. Proceedings of the National Academy of Sciences of the United States of America 101(38): 13909–13914. doi:10.1073/pnas.0404458101.

Farres, J., Martin-Caballero, J., Martinez, C., Lozano, J.J., Llacuna, L., Ampurdanes, C., et al. 2013. Parp-2 is required to maintain hematopoiesis following sublethal gammairradiation in mice. Blood 122(1): 44–54. doi:10.1182/blood-2012-12-472845.

Farres, J., Llacuna, L., Martin-Caballero, J., Martinez, C., Lozano, J.J., Ampurdanes, C., et al. 2015. PARP-2 sustains erythropoiesis in mice by limiting replicative stress in erythroid progenitors. Cell death and differentiation 22(7): 1144–1457. doi:10.1038/cdd.2014.202.

Fernandez-Garcia, B., Vaque, J.P., Herreros-Villanueva, M., Marques-Garcia, F., Castrillo, F., Fernandez-Medarde, A., et al. 2007. p73 cooperates with Ras in the activation of MAP kinase signaling cascade. Cell death and differentiation 14(2): 254–265. doi:10.1038/sj.cdd.4401945.

Frouin, I., Maga, G., Denegri, M., Riva, F., Savio, M., Spadari, S., et al. 2003. Human proliferating cell nuclear antigen, poly(ADP-ribose) polymerase-1, and p21waf1/cip1. A dynamic exchange of partners. The Journal of biological chemistry 278(41): 39265–39268. doi:10.1074/jbc.C300098200.

Fuertes, G., Martin De Llano, J.J., Villarroya, A., Rivett, A.J., and Knecht, E. 2003. Changes in the proteolytic activities of proteasomes and lysosomes in human fibroblasts produced by serum withdrawal, amino-acid deprivation and confluent conditions. The Biochemical journal 375(Pt 1): 75–86. doi:10.1042/BJ20030282.

Fujita, M., Watanabe, R., Jaensch, N., Romanova-Michaelides, M., Satoh, T., Kato, M., et al. 2011. Sorting of GPI-anchored proteins into ER exit sites by p24 proteins is dependent on remodeled GPI. The Journal of cell biology 194(1): 61–75. doi:10.1083/jcb.201012074.

Goldsmith, J.F., Lee, S.J., Jiang, J., and Frank, S.J. 1997. Growth hormone induces detergent insolubility of GH receptors in IM-9 cells. The American journal of physiology 273(5): e932. doi:10.1152/ajpendo.1997.273.5.E932.

Golenia, G., Gatie, M.I., and Kelly, G.M. 2017. Frizzled gene expression and negative regulation of canonical WNT-beta-catenin signaling in mouse F9 teratocarcinoma cells. Biochem. Cell. Biol. 95(2): 251–262. doi:10.1139/bcb-2016-0150.

Gungor-Ordueri, N.E., Sahin, Z., Sahin, P., and Celik-Ozenci, C. 2014. The expression pattern of PARP-1 and PARP-2 in the developing and adult mouse testis. Acta histochemica 116(5): 958–964. doi:10.1016/j.acthis.2014.03.010.

Gupte, R., Liu, Z., and Kraus, W.L. 2017. PARPs and ADP-ribosylation: recent advances linking molecular functions to biological outcomes. Genes Dev. 31(2): 101–126. doi:10.1101/gad.291518.116.

Hanzlikova, H., Gittens, W., Krejcikova, K., Zeng, Z., and Caldecott, K.W. 2017. Overlapping roles for PARP1 and PARP2 in the recruitment of endogenous XRCC1 and PNKP into oxidized chromatin. Nucleic Acids Res. 45(5): 2546–2557. doi:10.1093/nar/gkw1246.

Heiser, V., Scherzinger, E., Boeddrich, A., Nordhoff, E., Lurz, R., Schugardt, N., et al. 2000. Inhibition of huntingtin fibrillogenesis by specific antibodies and small molecules: implications for Huntington’s disease therapy. Proceedings of the National Academy of Sciences of the United States of America 97(12): 6739–6744. doi:10.1073/pnas.110138997.

Horigome, T., Furukawa, K., and Ishii, K. 2008. Purification and proteomic analysis of a nuclear-insoluble protein fraction. Methods Mol. Biol. 432: 139–148. doi:10.1007/978-1-59745-028-7_9.

Hottiger, M.O. 2015. Nuclear ADP-ribosylation and its role in chromatin plasticity, cell differentiation, and epigenetics. Annu. Rev. Biochem. 84: 227–263. doi:10.1146/annurev-biochem-060614-034506.

Huber, A., Bai, P., de Murcia, J.M., and de Murcia, G. 2004. PARP-1, PARP-2 and ATM in the DNA damage response: functional synergy in mouse development. DNA repair 3(8): 1103–1108. doi:10.1016/j.dnarep.2004.06.002.

Hwang, J.T., and Kelly, G.M. 2012. GATA6 and FOXA2 regulate Wnt6 expression during extraembryonic endoderm formation. Stem Cells Dev. 21(17): 3220–3232. doi:10.1089/scd.2011.0492.

Jeggo, P.A. 1998. DNA repair: PARP - another guardian angel? Current biology: CB 8(2): 49–51. doi:10.1016/S0960-9822(98)70032-6.

Jha, R., Agarwal, A., Mahfouz, R., Paasch, U., Grunewald, S., Sabanegh, E., et al. 2009. Determination of Poly (ADP-ribose) polymerase (PARP) homologues in human ejaculated sperm and its correlation with sperm maturation. Fertility and sterility 91(3): 782–790. doi:10.1016/j.fertnstert.2007.12.079.

Kamboj, A., Lu, P., Cossoy, M.B., Stobart, J.L., Dolhun, B.A., Kauppinen, T.M., et al. 2013. Poly(ADP-ribose) polymerase 2 contributes to neuroinflammation and neurological dysfunction in mouse experimental autoimmune encephalomyelitis. Journal of neuroinflammation 10: 49. doi:10.1186/1742-2094-10-49.

Kang, H.C., Lee, Y.I., Shin, J.H., Andrabi, S.A., Chi, Z., Gagne, J.P., et al. 2011. Iduna is a poly(ADP-ribose) (PAR)-dependent E3 ubiquitin ligase that regulates DNA damage. Proc. Natl. Acad. Sci. U S A 108(34): 14103–14108. doi:10.1073/pnas.1108799108.

Kashima, L., Idogawa, M., Mita, H., Shitashige, M., Yamada, T., Ogi, K., et al. 2012. CHFR protein regulates mitotic checkpoint by targeting PARP-1 protein for ubiquitination and degradation. The Journal of biological chemistry 287(16): 12975–12984. doi:10.1074/jbc.M111.321828.

Kawarabayashi, T., Younkin, L.H., Saido, T.C., Shoji, M., Ashe, K.H., and Younkin, S.G. 2001. Age-dependent changes in brain, CSF, and plasma amyloid (beta) protein in the Tg2576 transgenic mouse model of Alzheimer’s disease. J. Neurosci. 21(2): 372–381. doi:10.1523/JNEUROSCI.21-02-00372.2001.

Kilic, M., Schafer, R., Hoppe, J., and Kagerhuber, U. 2002. Formation of noncanonical high molecular weight caspase-3 and -6 complexes and activation of caspase-12 during serum starvation induced apoptosis in AKR-2B mouse fibroblasts. Cell Death Differ. 9(2): 125–137. doi:10.1038/sj.cdd.4400968.

Kraus, W.L. 2015. PARPs and ADP-ribosylation: 50 years … and counting. Molecular cell 58(6): 902–910. doi:10.1016/j.molcel.2015.06.006.

Krishnakumar, R., and Kraus, W.L. 2010. The PARP side of the nucleus: molecular actions, physiological outcomes, and clinical targets. Molecular cell 39(1): 8–24. doi:10.1016/j.molcel.2010.06.017.

Langelier, M.F., Riccio, A.A., and Pascal, J.M. 2014. PARP-2 and PARP-3 are selectively activated by 5′ phosphorylated DNA breaks through an allosteric regulatory mechanism shared with PARP-1. Nucleic acids research 42(12): 7762–7775. doi:10.1093/nar/gku474.

Ledesma, M.D., Bonay, P., Colaco, C., and Avila, J. 1994. Analysis of microtubule-associated protein tau glycation in paired helical filaments. The Journal of biological chemistry 269(34): 21614–21619.

Leung, A.K. 2014. Poly(ADP-ribose): an organizer of cellular architecture. The Journal of cell biology 205(5): 613–619. doi:10.1083/jcb.201402114.

Li, X., Klaus, J.A., Zhang, J., Xu, Z., Kibler, K.K., Andrabi, S.A., et al. 2010. Contributions of poly(ADP-ribose) polymerase-1 and -2 to nuclear translocation of apoptosis-inducing factor and injury from focal cerebral ischemia. J. Neurochem. 113(4): 1012–1022. doi:10.1111/j.1471-4159.2010.06667.x.

Lin, S.X., Ferro, K.L., and Collins, C.A. 1994. Cytoplasmic dynein undergoes intracellular redistribution concomitant with phosphorylation of the heavy chain in response to serum starvation and okadaic acid. The Journal of cell biology 127(4): 1009–1019. doi:10.1083/jcb.127.4.1009.

Liu, C., Wu, J., Paudyal, S.C., You, Z., and Yu, X. 2013. CHFR is important for the first wave of ubiquitination at DNA damage sites. Nucleic acids research 41(3): 1698–1710. doi:10.1093/nar/gks1278.

Lu, C., Shi, Y., Wang, Z., Song, Z., Zhu, M., Cai, Q., et al. 2008. Serum starvation induces H2AX phosphorylation to regulate apoptosis via p38 MAPK pathway. FEBS letters 582(18): 2703–2708. doi:10.1016/j.febslet.2008.06.051.

Maeda, Y., Hunter, T.C., Loudy, D.E., Dave, V., Schreiber, V., and Whitsett, J.A. 2006. PARP-2 interacts with TTF-1 and regulates expression of surfactant protein-B. The Journal of biological chemistry 281(14): 9600–9606. doi:10.1074/jbc.M510435200.

Martin, N., Schwamborn, K., Schreiber, V., Werner, A., Guillier, C., Zhang, X.D., et al. 2009. PARP-1 transcriptional activity is regulated by sumoylation upon heat shock. The EMBO journal 28(22): 3534–3548. doi:10.1038/emboj.2009.279.

Masdehors, P., Glaisner, S., Maciorowski, Z., Magdelenat, H., and Delic, J. 2000. Ubiquitin-dependent protein processing controls radiation-induced apoptosis through the N-end rule pathway. Experimental cell research 257(1): 48–57. doi:10.1006/excr.2000.4870.

Menissier de Murcia, J., Ricoul, M., Tartier, L., Niedergang, C., Huber, A., Dantzer, F., et al. 2003. Functional interaction between PARP-1 and PARP-2 in chromosome stability and embryonic development in mouse. The EMBO journal 22(9): 2255–2263. doi:10.1093/emboj/cdg206.

Mogi, M., Ozeki, N., Nakamura, H., and Togari, A. 2004. Dual roles for NF-kappaB activation in osteoblastic cells by serum deprivation: osteoblastic apoptosis and cell-cycle arrest. Bone 35(2): 507–516. doi:10.1016/j.bone.2004.03.003.

Nakashima, K., Yamazaki, M., and Abe, H. 2005. Effects of serum deprivation on expression of proteolytic-related genes in chick myotube cultures. Bioscience, biotechnology, and biochemistry 69(3): 623–627. doi:10.1271/bbb.69.623.

Nicolas, L., Martinez, C., Baro, C., Rodriguez, M., Baroja-Mazo, A., Sole, F., et al. 2010. Loss of poly(ADP-ribose) polymerase-2 leads to rapid development of spontaneous T-cell lymphomas in p53-deficient mice. Oncogene 29(19): 2877–2883. doi:10.1038/onc.2010.11.

Okano, S., Lan, L., Caldecott, K.W., Mori, T., and Yasui, A. 2003. Spatial and temporal cellular responses to single-strand breaks in human cells. Mol. Cell. Biol. 23(11): 3974–3981. doi:10.1128/MCB.23.11.3974-3981.2003.

Oliver, A.W., Ame, J.C., Roe, S.M., Good, V., de Murcia, G., and Pearl, L.H. 2004. Crystal structure of the catalytic fragment of murine poly(ADP-ribose) polymerase-2. Nucleic acids research 32(2): 456–464. doi:10.1093/nar/gkh215.

Paladino, S., Sarnataro, D., and Zurzolo, C. 2002. Detergent-resistant membrane microdomains and apical sorting of GPI-anchored proteins in polarized epithelial cells. International journal of medical microbiology: IJMM 291(6-7): 439–445. doi:10.1078/1438-4221-00151.

Peters, T.W., Rardin, M.J., Czerwieniec, G., Evani, U.S., Reis-Rodrigues, P., Lithgow, G.J., et al. 2012. Tor1 regulates protein solubility in Saccharomyces cerevisiae. Molecular biology of the cell 23(24): 4679–4688. doi:10.1091/mbc.E12-08-0620.

Puchi, M., Garcia-Huidobro, J., Cordova, C., Aguilar, R., Dufey, E., Imschenetzky, M., et al. 2010. A new nuclear protease with cathepsin L properties is present in HeLa and Caco-2 cells. Journal of cellular biochemistry 111(5): 1099–1106. doi:10.1002/jcb.22712.

Quenet, D., Gasser, V., Fouillen, L., Cammas, F., Sanglier-Cianferani, S., Losson, R., et al. 2008. The histone subcode: poly(ADP-ribose) polymerase-1 (Parp-1) and Parp-2 control cell differentiation by regulating the transcriptional intermediary factor TIF1beta and the heterochromatin protein HP1alpha. FASEB J 22(11): 3853–3865. doi:10.1096/fj.08-113464.

Refolo, L.M., Wittenberg, I.S., Friedrich, V.L., Jr., and Robakis, N.K. 1991. The Alzheimer amyloid precursor is associated with the detergent-insoluble cytoskeleton. J. Neurosci. 11(12): 3888–3897. doi:10.1523/JNEUROSCI.11-12-03888.1991.

Reis-Rodrigues, P., Czerwieniec, G., Peters, T.W., Evani, U.S., Alavez, S., Gaman, E.A., et al. 2012. Proteomic analysis of age-dependent changes in protein solubility identifies genes that modulate lifespan. Aging cell 11(1): 120–127. doi:10.1111/j.1474-9726.2011.00765.x.

Riccio, A.A., Cingolani, G., and Pascal, J.M. 2016. PARP-2 domain requirements for DNA damage-dependent activation and localization to sites of DNA damage. Nucleic acids research 44(4): 1691–1702. doi:10.1093/nar/gkv1376.

Rygiel, T.P., Mertens, A.E., Strumane, K., van der Kammen, R., and Collard, J.G. 2008. The Rac activator Tiam1 prevents keratinocyte apoptosis by controlling ROS-mediated ERK phosphorylation. Journal of cell science 121(8): 1183–1192. doi:10.1242/jcs.017194.

Schamberger, C.J., Gerner, C., and Cerni, C. 2005. Caspase-9 plays a marginal role in serum starvation-induced apoptosis. Experimental cell research 302(1): 115–128. doi:10.1016/j.yexcr.2004.08.026.

Scherzinger, E., Sittler, A., Schweiger, K., Heiser, V., Lurz, R., Hasenbank, R., et al. 1999. Self-assembly of polyglutamine-containing huntingtin fragments into amyloid-like fibrils: implications for Huntington’s disease pathology. Proceedings of the National Academy of Sciences of the United States of America 96(8): 4604–4609. doi:10.1073/pnas.96.8.4604.

Schreiber, V., Ame, J.C., Dolle, P., Schultz, I., Rinaldi, B., Fraulob, V., et al. 2002. Poly(ADP-ribose) polymerase-2 (PARP-2) is required for efficient base excision DNA repair in association with PARP-1 and XRCC1. The Journal of biological chemistry 277(25): 23028–23036. doi:10.1074/jbc.M202390200.

Shi, Y., Felley-Bosco, E., Marti, T.M., Orlowski, K., Pruschy, M., and Stahel, R.A. 2012. Starvation-induced activation of ATM/Chk2/p53 signaling sensitizes cancer cells to cisplatin. BMC cancer 12: 571. doi:10.1186/1471-2407-12-571.

Simmons, G., Gosalia, D.N., Rennekamp, A.J., Reeves, J.D., Diamond, S.L., and Bates, P. 2005. Inhibitors of cathepsin L prevent severe acute respiratory syndrome coronavirus entry. Proceedings of the National Academy of Sciences of the United States of America 102(33): 11876–11881. doi:10.1073/pnas.0505577102.

Singh, A., Upadhyay, V., Upadhyay, A.K., Singh, S.M., and Panda, A.K. 2015. Protein recovery from inclusion bodies of Escherichia coli using mild solubilization process. Microb. Cell. Fact. 14: 41. doi:10.1186/s12934-015-0222-8.

Takata, H., Nishijima, H., Ogura, S., Sakaguchi, T., Bubulya, P.A., Mochizuki, T., et al. 2009. Proteome analysis of human nuclear insoluble fractions. Genes to cells: devoted to molecular & cellular mechanisms 14(8): 975–990. doi:10.1111/j.1365-2443.2009.01324.x.

Tramontano, F., Malanga, M., and Quesada, P. 2007. Differential contribution of poly(ADP-ribose)polymerase-1 and -2 (PARP-1 and -2) to the poly(ADP-ribosyl)ation reaction in rat primary spermatocytes. Molecular human reproduction 13(11): 821–828. doi:10.1093/molehr/gam062.

Vida, A., Marton, J., Miko, E., and Bai, P. 2017. Metabolic roles of poly(ADP-ribose) polymerases. Semin Cell Dev Biol 63: 135–143. doi:10.1016/j.semcdb.2016.12.009.

Vidakovic, M., Grdovic, N., Quesada, P., Bode, J., and Poznanovic, G. 2004. Poly(ADP-ribose) polymerase-1: association with nuclear lamins in rodent liver cells. Journal of cellular biochemistry 93(6): 1155–1168. doi:10.1002/jcb.20289.

Waelter, S., Boeddrich, A., Lurz, R., Scherzinger, E., Lueder, G., Lehrach, H., et al. 2001. Accumulation of mutant huntingtin fragments in aggresome-like inclusion bodies as a result of insufficient protein degradation. Molecular biology of the cell 12(5): 1393–1407. doi:10.1091/mbc.12.5.1393.

Wang, T., Simbulan-Rosenthal, C.M., Smulson, M.E., Chock, P.B., and Yang, D.C. 2008. Polyubiquitylation of PARP-1 through ubiquitin K48 is modulated by activated DNA, NAD+, and dipeptides. Journal of cellular biochemistry 104(1): 318–328. doi:10.1002/jcb.21624.

Wei, H., and Yu, X. 2016. Functions of pARylation in DNA damage repair pathways. Genomics. Proteomics. Bioinformatics. 14(3): 131–139. doi:10.1016/j.gpb.2016.05.001.

Wong, S.L., Chan, W.M., and Chan, H.Y. 2008. Sodium dodecyl sulfate-insoluble oligomers are involved in polyglutamine degeneration. FASEB. J. 22(9): 3348–3357. doi:10.1096/fj.07-103887.

Wyrsch, P., Blenn, C., Bader, J., and Althaus, F.R. 2012. Cell death and autophagy under oxidative stress: roles of poly(ADP-Ribose) polymerases and Ca(2+). Mol. Cell. Biol. 32(17): 3541–3553. doi:10.1128/MCB.00437-12.

Xu, T.R., Lu, R.F., Romano, D., Pitt, A., Houslay, M.D., Milligan, G., et al. 2012. Eukaryotic translation initiation factor 3, subunit a, regulates the extracellular signal-regulated kinase pathway. Mol. Cell. Biol. 32(1): 88–95. doi:10.1128/MCB.05770-11.

Yelamos, J., Monreal, Y., Saenz, L., Aguado, E., Schreiber, V., Mota, R., et al. 2006. PARP-2 deficiency affects the survival of CD4+CD8+ double-positive thymocytes. The EMBO journal 25(18): 4350–4360. doi:10.1038/sj.emboj.7601301.

